# A quantitative theory for genomic offset statistics

**DOI:** 10.1101/2023.01.02.522469

**Authors:** Clément Gain, Bénédicte Rhoné, Philippe Cubry, Israfel Salazar, Florence Forbes, Yves Vigouroux, Flora Jay, Olivier François

## Abstract

Genomic offset statistics predict the maladaptation of populations to rapid habitat alteration based on association of genotypes with environmental variation. Despite substantial evidence for empirical validity, genomic offset statistics have well-identified limitations, and lack a theory that would facilitate interpretations of predicted values. Here, we clarified the theoretical relationships between genomic offset statistics and unobserved fitness traits controlled by environmentally selected loci, and proposed a geometric measure to predict fitness after rapid change in local environment. he predictions of our theory were verified in computer simulations and in empirical data on African pearl millet (*Cenchrus americanus*) obtained from a common garden experiment. Our results proposed a unified perspective on genomic offset statistics, and provided a theoretical foundation necessary when considering their potential application in conservation management in the face of environmental change.

## Introduction

### Maladaptation across environmental changes

Predicting maladaptation resulting from traits that evolved in one environment being placed in an altered environment is a long-standing question in ecology and evolution, originally termed as evolutionary traps or mismatches (Schlaepfer *et al*., 2002; Cook and Saccheri, 2013). With the increasing availability of genomic data, a recent objective is to determine whether those shifts could be predicted from the genetic loci that control adaptive traits and the fitness effects of these loci in spatially varying environments, bypassing any direct phenotypic measurements (Capblancq *et al*., 2020; Waldvogel *et al*., 2020). This question is crucial to understand whether sudden changes in the species ecological niche, *i.e.*, the sum of the habitat conditions that allow individuals to survive and reproduce, can be sustained by natural populations (Grinnell, 1917; Hutchinson, 1957; Sork *et al*., 2010; Jay *et al*., 2012; Aitken and Whitlock, 2013; Schoville *et al*., 2012; Foden *et al*., 2019). To this aim, several approaches have incorporated genomic information on local adaptation into predictive measures of population maladaptation across ecological changes, called genomic offset (or genomic vulnerability) statistics (Fitzpatrick and Keller, 2015; Capblancq *et al*., 2020; Waldvogel *et al*., 2020).

### Genomic offset statistics and their limitations

Genomic offset statistics first estimate a statistical relationship between environmental gradients and allelic frequencies using genotype-environment association (GEA) models (Forester *et al*., 2018). The inferred relationship is then used to evaluate differences in predicted allelic frequencies at pairs of points in the ecological niche (Fitzpatrick and Keller, 2015; Rellstab *et al*., 2016; Gougherty *et al*., 2021). The central hypothesis is that those statistics are predictive of changes in fitness traits that occur under altered environmental conditions (Capblancq *et al*., 2020). Recent efforts combining trait measurements in common garden experiments or natural population censuses with landscape genomic data have shown that the loss of fitness due to abrupt environmental shift correlates well with genomic offset predictions (Bay *et al*., 2018; Ruegg *et al*., 2018; Rhoné *et al*., 2020; Ingvarsson and Bernhardsson, 2020; Fitzpatrick *et al*., 2021; Chen *et al*., 2022; Sang *et al*., 2022). Experiments in which organisms are placed into an environment that differs from the one in which the traits evolved are, however, not always feasible (or efficient). Genomic offsets – that can be calculated in field studies – offer then a reasonable alternative to common garden experiments in a wide spectrum of applications to model and non-model organisms.

Despite substantial evidence for empirical validity, the proposed measures of genomic offset have well-identified limitations due to migration and gene flow (but see Gougherty *et al*. (2021)), population structure or genomic load. They also have difficulties to account for polygenic effects or correlated predictors (Rellstab *et al*., 2021; Aguirre-Liguori *et al*., 2021; Hoffmann *et al*., 2021). More importantly, different types of genomic offset statistics have been proposed in recent years (Fitzpatrick and Keller, 2015; Rellstab *et al*., 2016; Capblancq and Forester, 2021), and the inferred values for each of those statistics have not been explicitly linked to fundamental measures in quantitative and population genetics. The proposed measures lack theoretical foundations that would clarify how those different statistics are related to fitness and to each other. Thus, there is an urgent need to propose theoretical developments that will facilitate biological interpretations of genomic offset statistics. Here, we developed a theoretical framework that links genomic offset statistics to adaptive trait values controlled by ecological conditions, unifies existing approaches and addresses their limitations.

## Results

### Geometry of the ecological niche

We developed a geometric approach to the concept of genomic offset (GO) by defining a dot product of ecological predictors built on effect sizes of those predictors on allelic frequencies. Effect sizes, (**b**_ℓ_) = (*b*_ℓj_), were obtained from a GEA model of centered allelic frequencies on scaled predictors observed at a set of sampling locations. In that notation, *ℓ* stands for a locus, and *j* stands for a predictor. Effect sizes were corrected for the confounding effects of population structure and missing predictors (Methods:“GEA studies”). Given *d* ecological predictors, recorded in vector **x**, and their altered versions based on some change in time or space, recorded in **x**^*^, we defined a geometric GO – implemented as *genetic gap* in the computer package LEA – as a quadratic distance between the two vectors **x** and **x**^*^

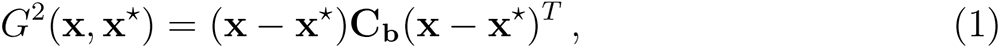

where 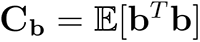 is the empirical covariance matrix of environmental effect sizes. Here the notation 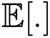 stands for the empirical mean across genomic loci in the analysis, ideally the number of loci controlling adaptive traits. Because the reference allele defining the genotype at a particular locus can be changed without any impact on the GEA analysis, we assume that the average value of effect sizes across all genomic loci is null, 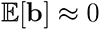. Considering allelic frequencies predicted from the GEA model, 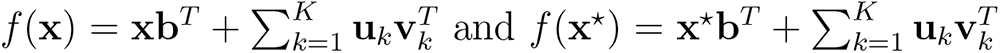, where the **u**_k_ represents

*K* confounding factors and **v**_k_ their loadings, we have

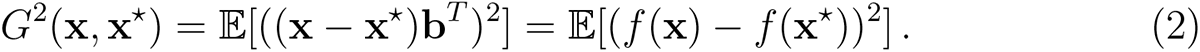

Thus the geometric GO has a dual interpretation as a quadratic distance in environmental and in genetic space. The population genetic interpretation of the geometric GO is as the average value of Nei’s *D*_ST_*/*2 (= *F*_ST_ *× H*_T_*/*2) for the set of loci assumed to be involved in local adaptation (Nei, 1973; François and Gain, 2021). As a genomic offset, the *D*_ST_ statistic can be calculated between pairs of population in space, but also in time, and it evaluates the genetic diversity between the populations in which **x** and **x**^*^ are measured or forecasted.

### Quantitative theory for genomic offset

We developed a quantitative theory for the geometric GO and for other GO statistics under the hypothesis of local stabilizing selection (Kimura, 1965; Lande, 1975). Under this hypothesis, observed allelic frequencies have reached local equilibria in which polygenic or quantitative characters are under natural selection for intermediate optimum phenotypes. The theory relies on a statistical model for an unobserved fitness trait for which a large number of small allelic effects mediate the effects of ecological predictors on fitness.

We defined *ω*(**x**, **x**^*^) to be the relative fitness value of a trait at equilibrium in environment **x** being placed in the altered environment **x**^*^. Under local Gaussian stabilizing selection, we found that the value of the logarithm of altered fitness varies in proportion with the geometric GO (Figure 1, Box 1)

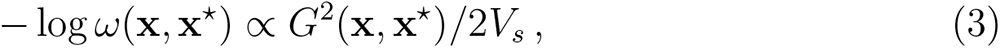

**Figure 1.**
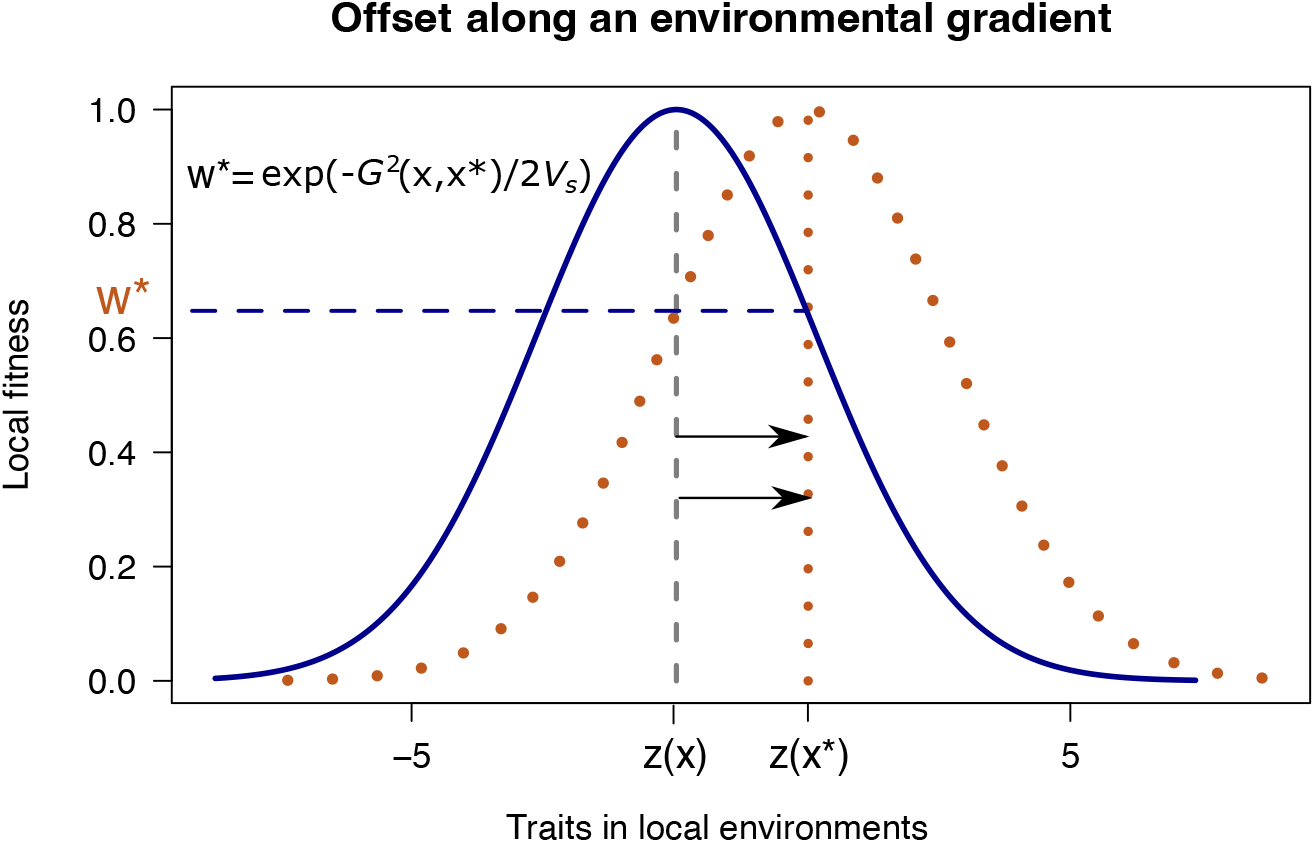
Geometric offset (genetic gap) under local Gaussian stabilizing selection. The two points, 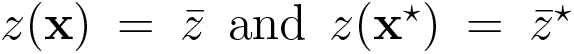, represent locally optimal values of an adaptive trait in respective environments **x** and **x**^*^. The curves display the fitness values for the trait in each environment. An organism with trait *z*(**x**), optimal in environment **x**, being placed in altered environment **x**^*^, has a fitness value equal to *ω*^*^ = exp(*−G*^2^(**x**, **x**^*^)*/*2*V*_s_), where *G*^2^(**x**, **x**^*^) is the genomic offset (horizontal dashed line), and *V*_s_ is defined in text.

where the *V*_s_ coefficient depends on the inherited variance and on the strength of stabilizing selection. In addition, the above equation remains valid when environmental predictors are indirectly related to the factors that influence the traits under selection, for example when those predictors are built on linear combinations of causal predictors for selection (Supporting Information: “Linear combination of predictors”). The geometric GO is thus robust to correlation in causal effects, and Eq. (3) extends to known and unknown linear combinations of those effects.

### Unifying genomic offset statistics

Beyond defining a new geometric measure of genomic offset, the quantitative theory provides a unified framework for GO statistics based on redundancy analysis (RDA, Capblancq and Forester (2021)), the risk of nonadaptedness (Rona, Rellstab *et al*. (2016)), and gradient forests (GF, Fitzpatrick and Keller (2015)) (Supporting Information:“Relationships to other GO statistics”). The main result is that all GO statistics predict the logarithm of fitness, but not for the same shape of the (within-locality) selection gradient. When RDA is performed on both environmental and latent predictors, the RDA GO is theoretically equivalent to the geometric GO, and thus predicts relative fitness under the hypothesis of Gaussian selection within localities. The risk of nonadaptedness, which is defined as the average of allelic frequency differences instead of squared differences, makes the implicit assumption that the selection gradient is built upon an exponential (Laplace) curve. When the distribution of effect sizes is Gaussian, Rona is then related to the square root of the geometric GO (times 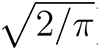). Like most machine learning techniques, GF is a nonparametric approach. In GF, no selection gradient is modelled a priori, but may be thought of as being estimated from the observed data. This might be one reason for which GF require more information than linear approaches based on low-dimensional parameters. The GF GO nevertheless follows a construction similar to the geometric GO and the RDA GO.

#### Box 1.

**Genomic offset theory**

Consider an (unobserved) fitness trait, *z*, for which a large number of genes mediate the effects of ecological predictors on organismal viability. Using Eq. (7) in (Barton *et al*., 2017), the trait value is assumed to be controlled*_√_*by *L* mutations each having infinitesimally small allelic effect of equal size, 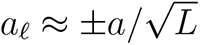, defining the trait value as a polygenic score, 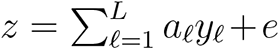. Here, *y*_ℓ_ is the allelic frequency at locus *ℓ*, expressed as deviation from the population mean, *a*_ℓ_ has random sign, *a*^2^ controls the additive genetic variance, and the random term *e* models the non-genetic variance. The definition is equivalent to the more traditional decomposition of variance into inherited and non-inherited components (Figure S1). Assuming a local Gaussian stabilizing selection model, the relative fitness of the trait in environment **x** is equal to 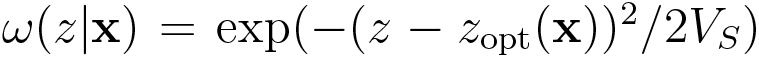, where 1/*V_s_* represents the strength of stabilizing selection. Conditional on local environment, the optimum, *z*_opt_(**x**), corresponds to the mean (or predicted) value of the trait, 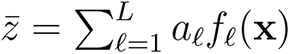. The logarithm of fitness for a trait at equilibrium in environment **x** being placed in the altered environment **x**^*^ is thus equal to

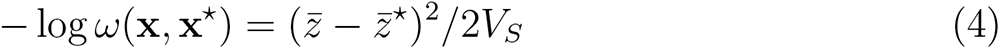

where 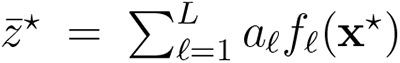. The difference in fitness traits 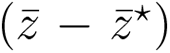, is equal to 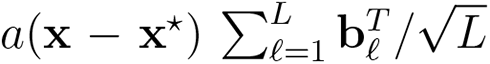. According to the central limit theorem, the conditional distribution of 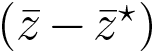 is Gaussian 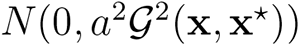, where 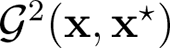 is defined from the theoretical – instead of empirical – effect size covariance matrix. The distribution of 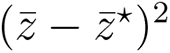 is a non-standard chi-squared distribution with one degree of freedom

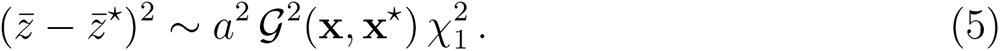

Since 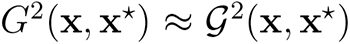 for large *L*, the value of the logarithm of altered fitness varies in proportion with the geometric GO, where the proportionality coefficient is equal to 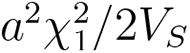. The expected value is thus approximately equal to 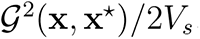, where *V*_s_ = *V*_S_*/a*^2^. Consideration of traits that are not at equilibrium in environment **x** adds an intercept term to the expected value, equal to 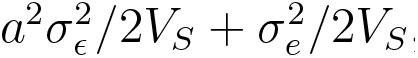 where 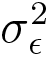 is the residual variance in the GEA model and 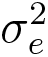 is the non-inherited variance (Supporting Information: “Logarithm of altered fitness for non-optimal traits”).

### Validation of the theory

To illustrate the above theory, we analyzed simulated data in which adaptive traits were matched to ecological gradients by local Gaussian stabilizing selection (Figure 2A, Methods:“Simulation study”, Supporting Information:“Extended simulation study”) (Haller and Messer, 2019). Two environmental predictors playing the role of temperature and precipitation in the studied range were considered, as well as two additional non-causal predictors correlated to the first ones (Figure 2B). The median values of temperature and precipitation determined four broad types of environments from *dry/warm* to *wet/cold* conditions. As an outcome of the simulation, the genetic groups resulting from selection, drift and gene flow matched the environmental classes, generating high levels of correlation between environmental predictors and population structure in the GEA analysis (Figure S2). As predicted by equation (3), the values of the geometric GO computed according to equation (1) varied linearly with the logarithm of fitness after alteration of local conditions (*r*^2^ *≈* 78%*, P <* 0.001, Figure 2C-D). The predictive power of the geometric GO was much higher than the predictive power of squared Euclidean environmental distance between predictors and their altered values (*r*^2^ *≈* 45%, *J* = 11.3, *P <* 0.001). Although it was calculated on both causal and non-causal predictors, the GO adjusted almost perfectly to the quadratic function that determines the intensity of local Gaussian stabilizing selection (*r*^2^ = 97%, *P <* 0.001, Figure S3). The first two eigenvalues of the covariance matrix of environmental effect sizes were much larger than the last ones (Figure 2C). We found that the loadings on the first axes gave more weight to predictors associated with natural selection, while the loadings on the last axes weighted predictors that did not play a role in the simulated evolutionary process. Uninformative predictors were given only low weights in the calculation of the GO statistic. Those results provided evidence that the largest eigenvalues that characterize the geometric GO contain useful information about local adaptation.

**Figure 2.**
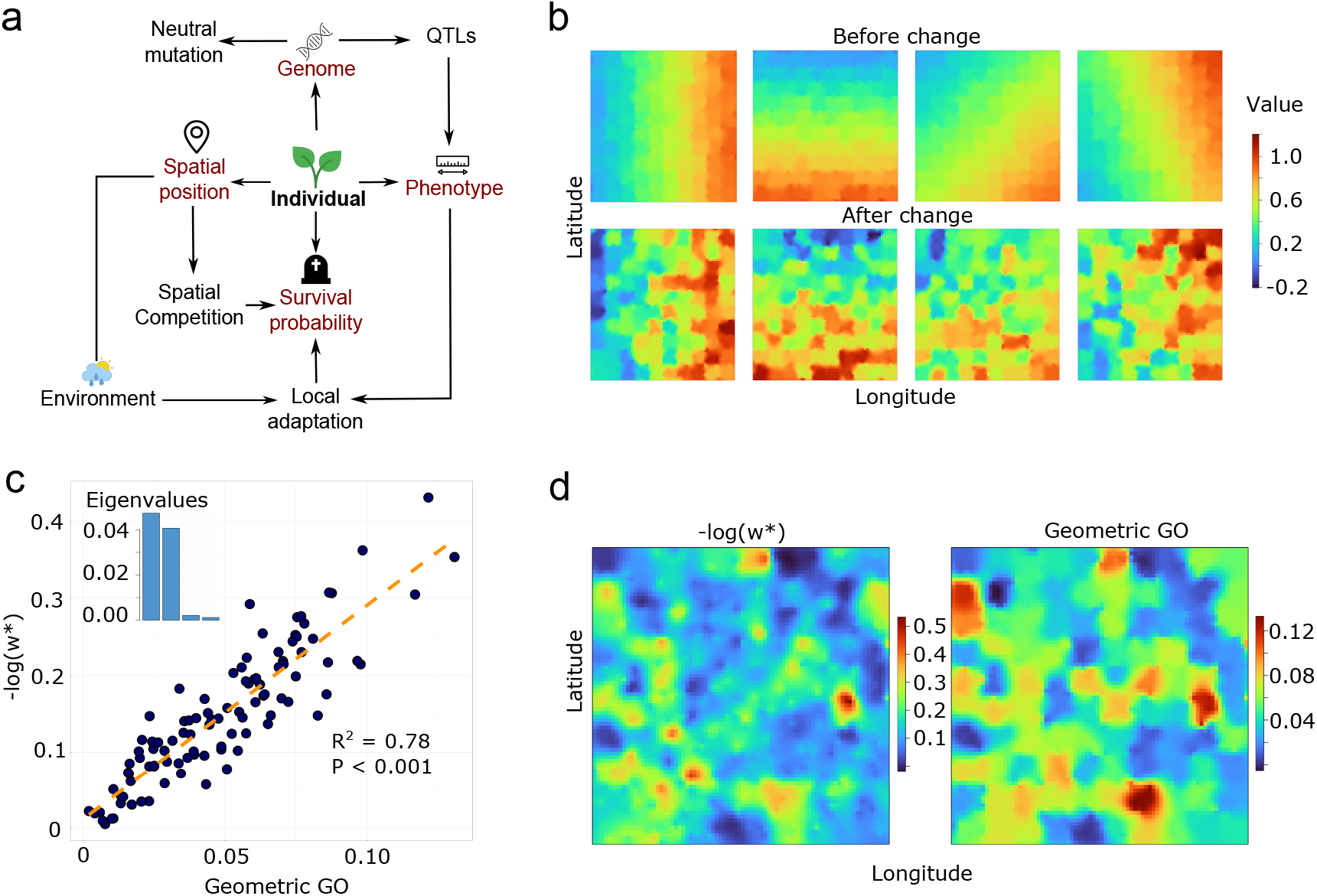
Simulation of fitness traits and geometric offset. **a**) Spatial individual-based forward simulations: Adaptive traits were matched to ecological gradients by local Gaussian stabilizing selection. **b**) Geographic maps of four environmental predictors before and after change. **c**) Logarithm of altered fitness values as a function of geometric offset. The eigenvalues of the covariance matrix of environmental effect sizes are displayed in the top left corner. **d**) Geographic maps of the logarithm of altered fitness values (left) and geometric offset (right).

### Extended simulation study

Expanding our case analysis, additional simulation scenarios were considered with traits under local stabilizing selection having distinct levels of polygenicity. Some cases were complicated by a strong correlation of environmental predictors with population structure. To overcome this complication, correction based on latent factors were included in all GO calculations (Methods:“GO computations”). As predicted by the theory, the values of the squared correlation between the GO statistic and the logarithm of fitness were very close to each other in all investigated cases (Figure 3, Figure S4). As expected, methods that did not use correction (undercorrection) or include population structure covariates (overcorrection) worked less well than methods with latent factor correction (Figures S5-S6). Once corrected, the GO statistics ranked similarly in all simulation scenarios. The ability of the geometric GO to predict the logarithm of fitness was equal to that of corrected RDA GO. It was slightly superior to that of Rona and to that of the GF GO. All GO statistics were highly correlated with the geometric GO (Figure S7). The geometric GO also exhibited high correlation with the quadratic distance between causal predictors explaining the traits under local stabilizing selection in the simulation model (Figure S8). This result supported the evidence of near-optimal fitness prediction by the GO statistics in all simulated evolutionary scenarios. When all genomic loci in the genotype matrix were included in the GO calculations, the predictions stayed close to those based on subsets of loci identified in the GEA analysis, GF GO reaching then performances similar to the other GO statistics (Figure S9).

**Figure 3.**
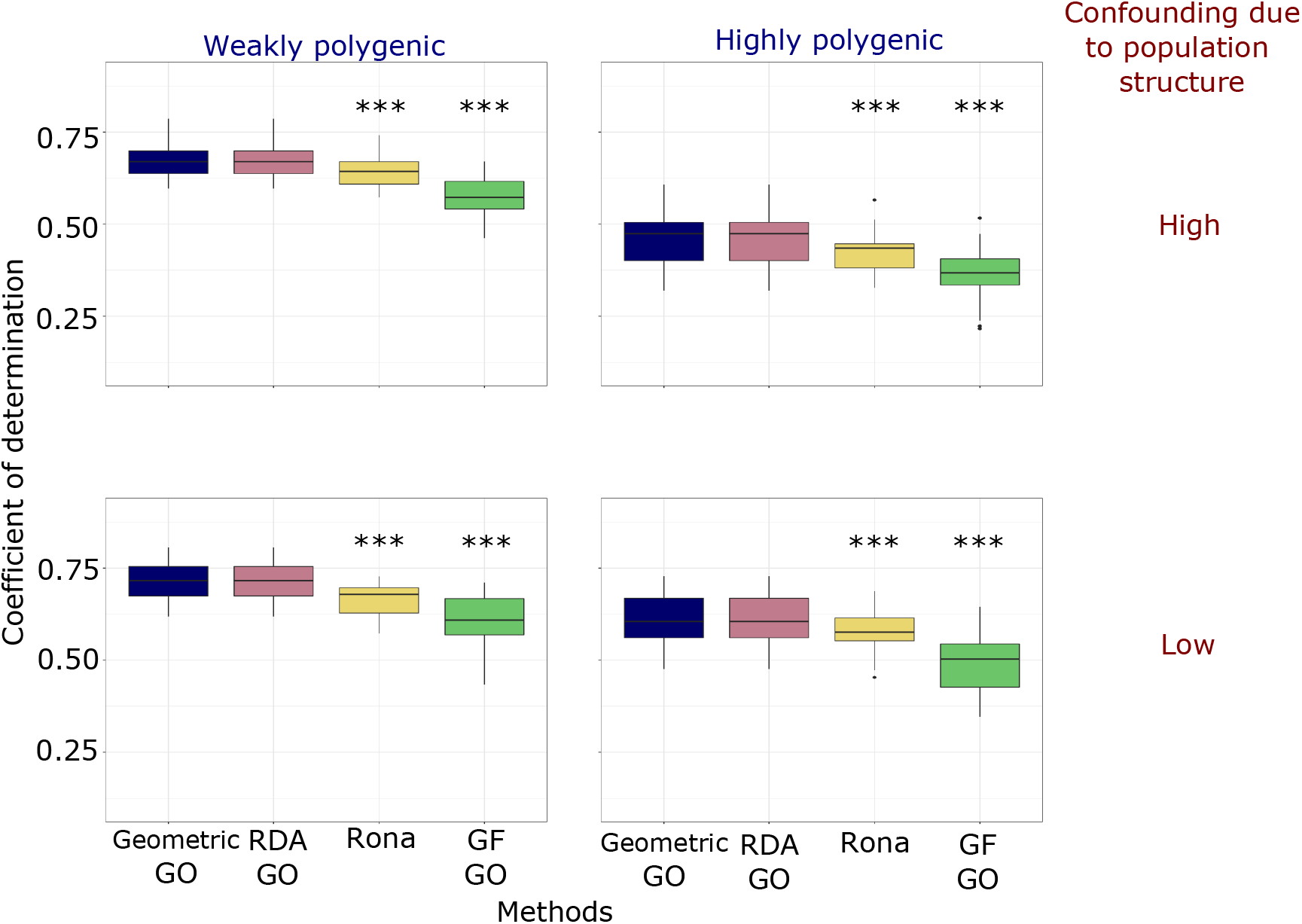
Predictive performances of GO statistics. Proportion of variance of fitness in the altered environment explained by GO statistics (coefficient of determination). Four scenarios with distinct levels of polygenicity in adaptive traits and correlation of environmental predictors with population structure were implemented. Significance values were based on paired t-tests of the difference in mean performance for each GO statistic relative to the geometric GO (***: *P <* 0.001). Boxplots display the median, the first quartile, the third quartile, and the whiskers of distributions. The upper whisker extends from the hinge to the largest value no further than 1.5 inter-quartile range (IQR) from the hinge. The lower whisker extends from the hinge to the smallest value at most 1.5 IQR of the hinge. Extreme values are represented by dots.

### Evaluating the bias of linear allelic frequency predictions

An approximation made by the geometric and other GO statistics is that allelic frequencies are predicted by unconstrained linear functions of environmental predictors. To evaluate the impact of this approximation, we compared linear predictions to those of a logistic regression model, which are constrained between zero and one. For small environmental change, the effect sizes in the linear GEA model could be approximated by the effect sizes in the logistic regression multiplied by the heterozygosity at each locus (Supporting Information:“Bias of linear predictors”). The geometric GO was then accurately approximated by the squared distance between constrained genetic predictors, E[(*f*_c_(**x**) *− f*_c_(**x**^*^))^2^] (Figure S10). Using a nonlinear machine learning model (Supporting Information:“Variational autoencoder GO”), we found again that the squared genetic distance between constrained genetic predictors strongly correlated with the geometric GO, supporting the approximation of fitness in altered environment using linear models (Figure S11).

### Pearl millet common garden experiment

We hypothesized that GO statistics could predict the logarithm of fitness in pearl millet, a nutritious staple cereal cultivated in arid soils in sub-Saharan Africa (Rhoné *et al*., 2020). Pearl millet is grown in a wide range of latitudes and climates with wide variety of ecotypes (landraces). The geometric GO and other measures of GO were estimated from 138,948 single-nucleotide polymorphisms for 170 Sahelian landraces in a two-year common garden experiment conducted in Sadoré (Niger) using loci identified in the GEA study (Figure 4A, Methods:“Pearl millet experiment”). For each landrace grown in the common garden, the total weight of seeds was measured as a proxy of landrace fitness, which was explained by a Gaussian selection gradient (Figure S12). Including latent factor correction, GO statistics were computed using the climate condition at the location of origin of the landrace and the climate at the experimental site. All GO statistics displayed a consistent relationship with the logarithm of seed weight (Figure 4B, Figure 5). Loci identified in the GEA study increased the performance of GO statistics compared to using whole genomic data, and the improvements were substantial compared to methods that did not include correction for confounding factors (Figures S13-S14 and Table S1). The best predictions of fitness in the common garden were obtained with the geometric GO and with the corrected version of Rona (*r*^2^ = 61%, *P <* 0.001, Figure 5). The eigenvalues and eigenvectors of the covariance matrix of environmental effect sizes suggested that climatic conditions could be summarized in three axes. Temperature predictors were given higher importance in driving fitness variation than precipitation and solar radiation predictors (Figure S15).

**Figure 4.**
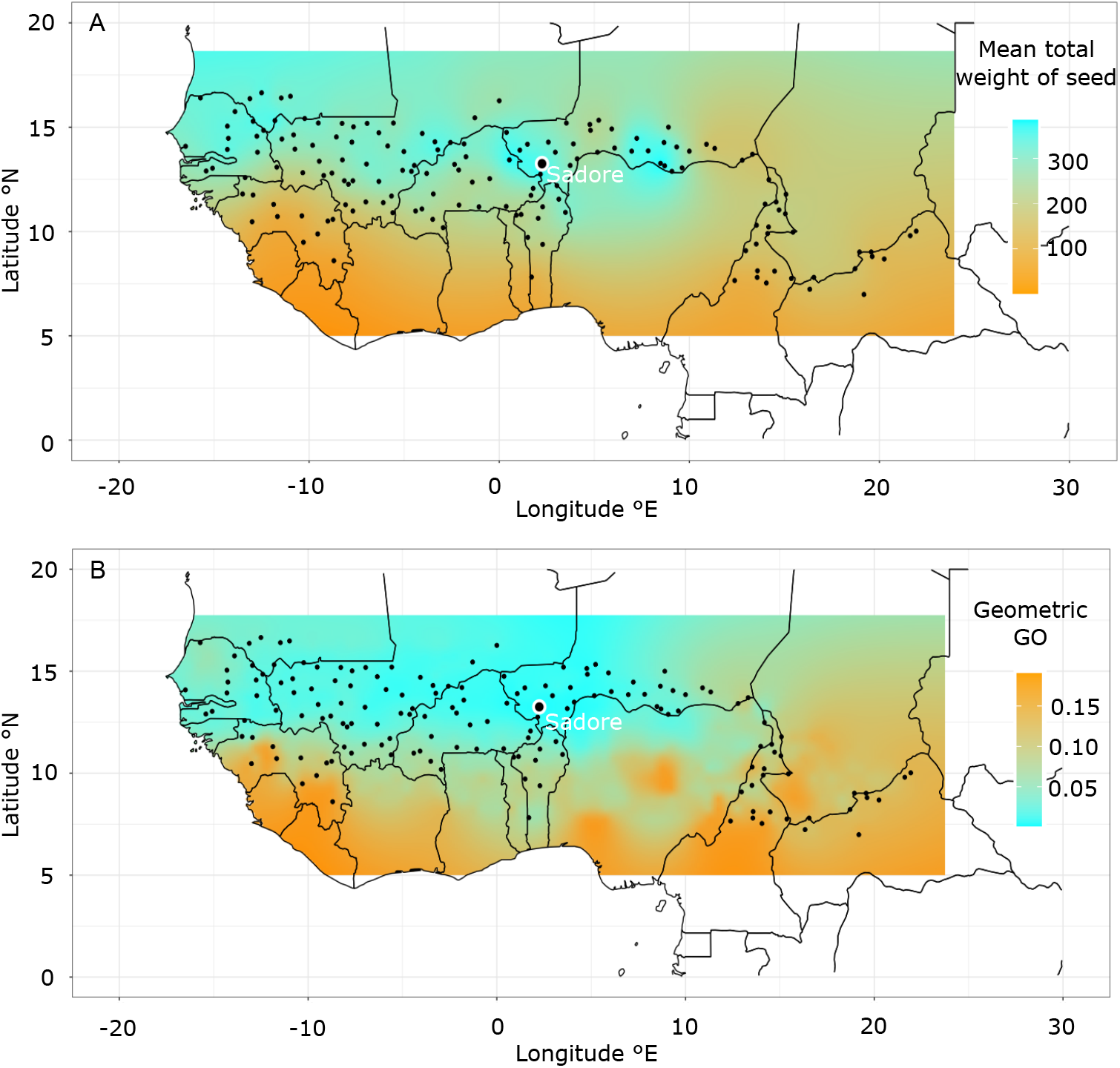
Interpolated fitness gradient and genomic offset for pearl millet landraces. A) Fitness values (log) measured as the mean total seed weight for each pearl millet landrace in the common garden experiment located in Sadoré (Niger). B) Values of the geometric genomic offset. Locations of landrace origin are represented as dots. Values at unsampled locations were interpolated from the nearest sampled location using the inverse distance weighting method.

**Figure 5.**
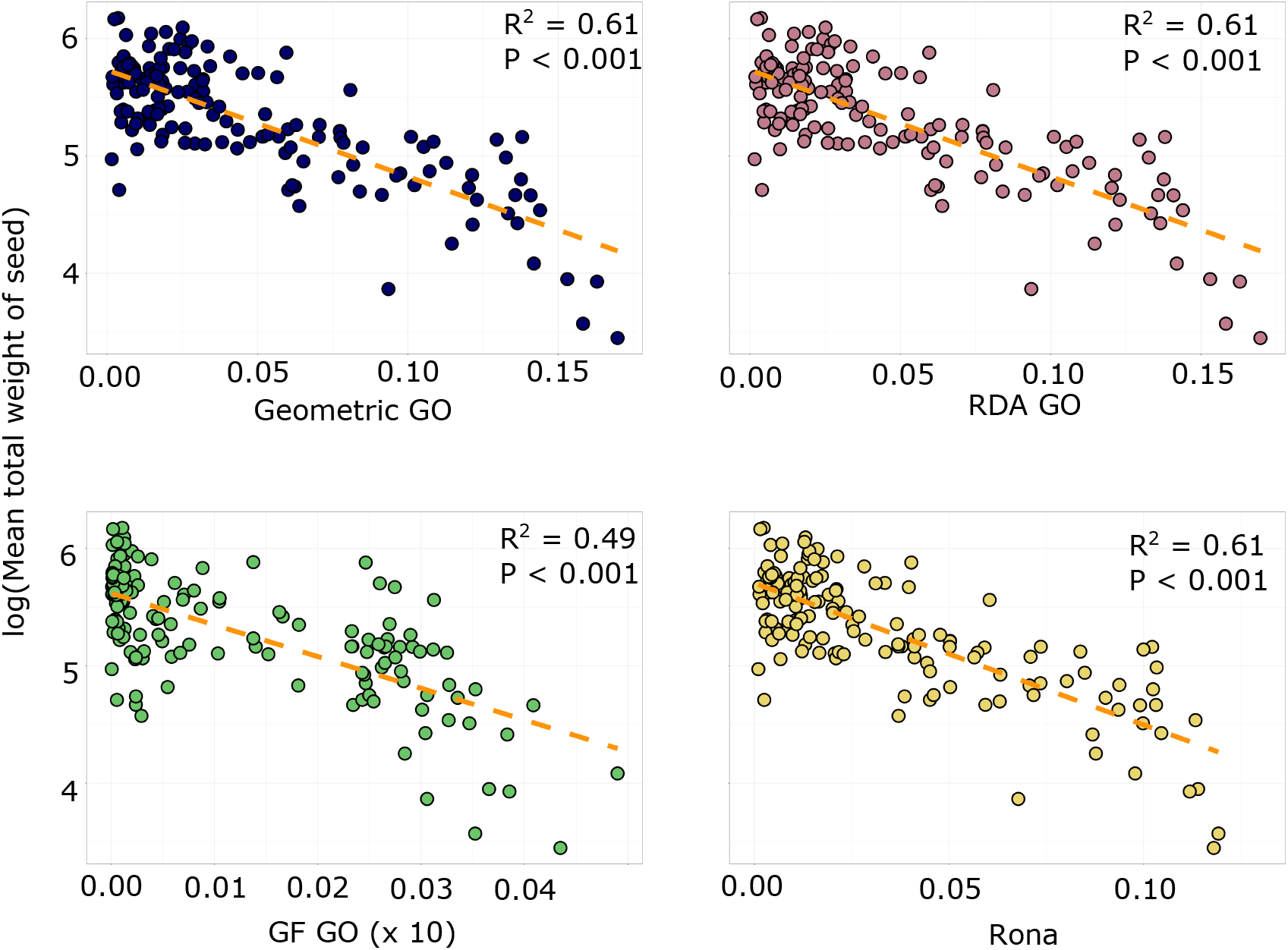
Logarithm of fitness in the common garden as a function of the GO statistic. Latent factor corrections were included in the calculation of all GO statistics (ten factors). Fitness was evaluated as the mean total weight of seed for 170 pearl millet landraces. GO values for GF were multiplied by a factor of ten.

## Discussion

### Quantitative theory

The geometric theory presented in our study provided a unified framework that not only explains why and when a GO statistic differs from the standard Euclidean environmental distance, but also allowed for a better understanding of previous measures of genomic offset. Based on models of local selection gradients, a theoretical analysis of GO statistics relying on Fisher’s infinitesimal trait model was developed. In this framework, the geometric GO decays linearly with the logarithm of fitness in the altered environment. Although of much lower computational complexity, the geometric GO was proved to be equivalent to a GO based on RDA, which justifies the use of RDA approaches under local Gaussian selection. The square root of the geometric GO was connected to Rona, and justifies the use of absolute differences in allele frequencies under exponential selection gradient.

### Improving GO statistics

According to Rellstab *et al*. (2021), current GO statistics may provide wrong predictions due to the correlation between population structure at selectively neutral loci and environmental predictors. Built on unbiased effect sizes, the geometric GO, which is based on a unique model for GEA estimation and for GO prediction, addressed this problem by including latent factors as covariates in the prediction model. Latent factor corrections were then incorporated into all considered GO statistics, which increased their predictive performance compared to their traditional usage. Our versions of RDA GO and Rona – that slightly differ from original proposals – were implemented in the R package LEA. Although those changes led to improved statistics, the geometric GO reached higher predictive performance than the other GO approaches. Next, the geometric GO addressed the problem of correlated predictors by modeling the covariance of their effect sizes. The importance of predictors could be assessed by examining the eigenvalues and eigenvectors of the environmental effect size covariance matrix. The eigenvalues provide a natural ranking of the importance of each axis, similar to the cumulative importance curves in GF. When a statistical analysis includes redundant predictors, reproducing information already present in a reduced set of predictors, the geometric GO gave lower weight to those redundant predictors, and differed substantially from the Euclidean environmental distance. Generally, the principal benefit of genomic offset over purely environmental distances in predicting maladaptation comes from the weighting of environmental predictors by their effect sizes (Làruson *et al*., 2022). All proposed GO approaches share the principle of weighting the environmental predictors by their strength of genetic association. For the vast majority of organisms where the most important predictors are unknown or for which common garden experiments are not efficient or unfeasible, genomic offset therefore provides a useful means for weighting the environmental predictors based on the information contained in allele frequencies.

### Limitations

Our simulation models and our theoretical developments relied upon a model of genotype *×* environment interaction for fitness related to antagonistic pleiotropy, whereby native alleles are best adapted to local conditions (Kawecki, 2004; Anderson *et al*., 2011). While antagonistic pleiotropy is an important mechanism for local adaptation, there are other types of interactions for fitness. If local adaptation is caused by conditional neutrality at many loci, where alleles show difference in fitness in one environment, but not in a contrasting environment, the predictive performances of GO statistics remain to be explored. In addition, GO statistics (except GF) are based on linear models for the relationship between genotype and environment. Linear models generate GO statistics that are invariant under translation in the niche, making predictions relevant at the center of the species distribution, but perhaps less relevant at margins of the range. While translational invariance could be corrected for by defining the offset as the average of squared differences between allelic frequencies in nonlinear models, we found that the results were very close to the linear models. An explanation may be that nonlinear machine learning models offer more flexible GO statistics than linear models, but that linear models achieve a better bias-variance trade-off than machine learning models, likely because less data are needed for their application. Other conceptual limitations include gene flow and constraints on adaptive plasticity that might mitigate the effect of environmental change on fitness (Aguirre-Liguori *et al*., 2021; Kawecki, 2004). As they do not use any observed information on fitness traits, GO statistics provide measures of expected fitness loss based on the indirect effects of environment mediated by loci under selection (Baron and Kenny, 1986). GO statistics are more accurate when non-genetic effects do not covary with environmental predictors. Lastly, we found that using candidate loci based on statistical significance in GEA improved prediction of fitness in altered conditions both in simulation and in real data analysis. We think that this happens because those studies may generally be underpowered, *i.e.*, a much larger sample size would increase the predictive power of GO statistics. Using a liberal threshold in GEA studies was considered as a trade-off between polygenicity and statistical significance, so that the GO measures could actually be based on polygenic scores while not erasing or blurring the genomic signals of local adaptation.

### Pearl millet experiment

To compare predictions of local adaptation with empirical data, GO statistics were estimated in a common garden experiment on pearl millet landraces in sub-Saharan Africa. Using GF, the original study reported a squared correlation of *r*^2^ *≈* 9.5 *−* 17% for seed weight, indicating that higher genomic vulnerability was associated with lower fitness under the climatic conditions at the experimental site (Rhoné *et al*., 2020). In our reanalysis, signals of local adaptation were consistent across all GO statistics, and improved fitness prediction substantially, up to a value of squared correlation equal to *r*^2^ *≈* 61%. The results strengthened the conclusions of (Rhoné *et al*., 2020), and supported the use of GO statistics in predictions of fitness values across the sub-Saharan area.

### Conclusions

Considering a duality between genetic space and environmental space, we developed a theoretical framework that linked GO statistics to a non-Euclidean geometry of the ecological niche. The geometric GO, as well as the modified Rona statistic, were implemented in the genetic gap function of the R package LEA (Gain and François, 2021). As a result of the quantitative theory, interpretations in terms of fitness in the altered environment were proposed, unifying several existing approaches, and addressing some of their limitations. Based on extensive numerical simulations and on data collected in a common garden experiment, our study indicated that GO statistics are important tools for conservation management in the face of climate change.

## Materials and Methods

### GEA studies

GEA studies and estimates of environmental effect sizes were performed based on LFMMs in the computer package LEA v3.9 (Caye *et al*., 2019; Gain and François, 2021). In LFMMs, allelic frequencies are modelled at each genomic locus of a genotype matrix as a mixed response of observed environmental variables with fixed effects and *K* unobserved latent factors. The number of latent factors was estimated from the screeplot of a principal component analysis of the genotype matrix. Loci with minor allele frequency less than 10% were filtered out the analysis. Statistical significance was determined by using the R package qvalue at a level of false discovery rate equal to 10%.

### GO computations

RDA was performed by using principal components of fitted values of the GEA regression model. Rona was computed as the average value of the absolute distance between predicted allelic frequencies across genomic loci (de Aquino *et al*., 2022; Rellstab *et al*., 2016). GF computations were performed using the R package gradientForest version 0.1. For consistency, we reported squared values of GO statistics in RDA and GF. Unless specified, GO statistics were computed on the loci detected in the GEA study, i.e., a same set of loci for all methods. To correct statistics for the confounding effect of population structure, all analyzes were performed conditional on the factors estimated in the LFMM analysis (Supporting Information:“GO computations”).

### Simulation study

Spatially-explicit individual-based simulations were performed using SLiM 3.7 (Haller and Messer, 2019) (Supporting Information:“Extended simulation study”). Each individual genome contained neutral mutations and quantitative trait loci (QTLs) under local stabilizing selection from a two-dimensional environment. The probability of survival of an individual genome in the next generation was computed as the product of density regulation and fitness. We designed four classes of scenarios, including weakly or highly polygenic traits, and weak or high correlation of environment with population structure. In scenarios with high polygenicity, traits controlled by 120 mutations with additive effects were matched to each environmental variable by local stabilizing selection. In weakly polygenic scenarios, the traits were controlled by 10 mutations. Scenarios with high confounding effects were initiated in a demographic range expansion process, creating correlation between environment and allelic frequencies at the genome level. For each scenario, thirty replicates were run with distinct seed values of the random generator. At the end of a simulation, individual geographic coordinates, environmental variables and individual fitness values before and after instantaneous environmental change were recorded. Paired *t*-tests were used to test statistical differences in the mean of predictive performances for the geometric GO and the other GO statistics.

### Empirical study

Methods regarding the common garden experiment on Pearl millet landraces conducted in Sadoré (13*^◦^* 14’ 0” N, 2*^◦^* 17’ 0” E, Niger, Africa) were described by Rhoné *et al*. (2020). For each of 170 landraces grown in the common garden, the total weight of seeds was measured by harvesting the main spike in ten plants per landrace sown during two consecutive years and was used as a proxy of landrace fitness. For each landrace grown in the common garden, environmental predictors, **x**, were obtained at the location of origin of the landrace, and **x**^*^ corresponded to the local conditions in Sadoré. We made the hypothesis that the mean total weight of seeds for a landrace was proportional to *ω*(**x**, **x**^*^) in the common garden. Using 100 plants per landrace in a pool-sequencing design, allelic frequencies were inferred at 138,948 single-nucleotide polymorphisms. Climate data were used to compute 157 metrics in three categories, precipitation, temperature (mean, maximum and minimum near surface air temperature), and surface downwelling shortwave radiation, that were reduced by principal component analysis (27 axes). GO statistics were computed using the climate condition (**x**) at the location of origin of the landrace and the climate conditions (**x**^*^) at the experimental site. For each GO statistic, we estimated a linear relationship with the logarithm of the mean total weight of seeds and used Pearson’s squared correlation to evaluate the goodness of fit. The *J* –test was used to test differences between predictive performances, corresponding to *R*squared for distinct regression models, of the geometric GO and other GO statistics (Davidson and MacKinnon, 1981).

## Data Availability

The pearl millet data have already been published, and have permissions appropriate for fully public release.

## Code Availability

The codes necessary to reproduce the simulations and data analyses of this study are available at https://github.com/bcm-uga/geneticgap under GNU General Public License v3.0 The geometric GO is implemented in the genetic gap function of the R package LEA (version number *>* 3.9.5) available from the public repository bioconductor and https://github.com/bcm-uga/LEA (latest version).

## Acknowledgments

This work received support from the French National Research Agency, projects PEG (grant number ANR-22-CE45-0033), Afradapt (grant number ANR-22-CE32-0008), and ETAPE (grant number ANR-18-CE36-0005). The authors are grateful to Thibaut Capblancq for many interactions and fruitful discussions.

## Author contributions

B.R., P.C., Y.V., I.S., F.F. contributed analyzes and helped drafting the manuscript. C.G., F.J. and O.F. conceived the study, developed the method, carried out analyzes, and wrote the manuscript.

## Additional information

Supporting methods and materials, supplementary figures and tables, are available online.

