## Supporting Information for "A quantitative theory for genomic offset statistics"

#### Contents

**1 Details on the geometric GO computation 2**

**2 Logarithm of altered fitness for non-optimal traits 4**

**3 Linear combination of predictors 5**

**4 Relationships between GO statistics 5**

**8 Code and data availability 21**

### 1 Details on the geometric GO computation

This section provides details on the mathematical definition of the geometric GO (genetic gap in computer implementation) introduced in the main text paragraph “Geometry in the ecological niche”. For a sample of $n$ genotypes, a GEA model adjusts the genotype matrix from a set of $d$ environmental predictors. Environmental predictors are scaled, so that they are unitless, and allelic frequencies are centered, so that their mean values across samples is zero. Environmental predictors and allelic frequency responses are thus expressed as deviations from population average. For $d$ environmental predictors considered as descriptors of the ecological niche, the GEA model, a latent factor mixed model (LFMM), estimates a matrix of effect sizes, $\mathbf{B} = (b_{\ell j}) = (\mathbf{b}_{\ell})$, with coefficients corresponding to the effect of each environmental predictor, $j = 1, \ldots, d$, on allelic frequency at each genetic locus, $\ell = 1, \ldots, L$, where $L$ is the number of loci in the genotype matrix (Caye *et al.*, 2019; Gain and François, 2021). The dimension of the $\mathbf{B}$ matrix is equal to $L \times d$, representing $L$ vectors in $d$ dimensions. For two environmental vectors $\mathbf{x}$ and $\mathbf{x}^*$ in the $d$-dimensional space, a dot product is defined as $\langle \mathbf{x}, \mathbf{x}^\star \rangle_{\mathbf{b}} = \mathbf{x}\mathbf{C}_{\mathbf{b}}\mathbf{x}^{\star T}$, where $\mathbf{C}_{\mathbf{b}} = \mathbf{B}^T\mathbf{B}/L = \mathbb{E}[\mathbf{b}^T\mathbf{b}]$ is the $d \times d$ empirical covariance matrix of effect sizes. More generally, the mathematical notation $\mathbb{E}[.]$ stands for an average over the $L$ genomic loci included in the analysis. If $\mathbf{q}_\ell$ is a quantity evaluated at locus $\ell$, then $\mathbb{E}[\mathbf{q}] = \sum_{\ell=1}^{L} \mathbf{q}_\ell / L$.

Based on this non-Euclidean geometry, the geometric GO is defined as the squared norm of the difference between two predictor vectors, $\mathbf{x}$ and $\mathbf{x}^\star$,

$$G^2(\mathbf{x}, \mathbf{x}^\star) = \langle \mathbf{x} - \mathbf{x}^\star, \mathbf{x} - \mathbf{x}^\star \rangle_{\mathbf{b}} = (\mathbf{x} - \mathbf{x}^\star)\mathbf{C}_{\mathbf{b}}(\mathbf{x} - \mathbf{x}^\star)^T = \|\mathbf{x} - \mathbf{x}^\star\|^2_{\mathbf{b}} .$$

To make a distinction between the empirical covariance and its theoretical value for a large number of loci, we denote by $\mathcal{G}^2(\mathbf{x}, \mathbf{x}^\star)$ the offset defined from the theoretical covariance matrix.

The square root of $G^2(\mathbf{x}, \mathbf{x}^\star)$ defines a mathematical distance between the environmental vectors $\mathbf{x}$ and $\mathbf{x}^\star$. In our empirical data, $\mathbf{x}^\star$ will represent environmental predictors in a common garden, and $\mathbf{x}$ will represent environmental predictors in the geographic location at each sample origin. The geometric GO extends to any pair of environmental predictors for current or future local environments, regardless of their geographic origin or their predicted value in time.

### 2 Logarithm of altered fitness for non-optimal traits

Consideration of fitness values for a trait optimal under environment $\mathbf{x}$ being placed in altered environment $\mathbf{x}^\star$, $\omega(\mathbf{x}, \mathbf{x}^\star) = \omega(z_{\text{opt}}(\mathbf{x})|\mathbf{x}^\star)$, can be viewed as a simplification for altered fitness in the case of near-optimal traits. The definition of altered fitness can be extended to near-optimal traits, $z$, by considering the random quantity $\omega(z|\mathbf{x}^\star)$.

According to the theory (Box 1 in main text), the trait value $z$ can be decomposed as $z = a(\mathbf{x}\mathbf{b}^T + \mathbf{u}\mathbf{v}^T + \epsilon) + e$, where $\mathbf{b} \sim \mathbf{N}(0, \mathbf{C}_\mathbf{b})$ , $\epsilon \sim N(0, \sigma_\epsilon^2)$ and $e \sim N(0, \sigma_e^2)$, after the central limit theorem. The first term corresponds to the additive genetic variance mediated by ecological predictors. The second term corresponds to the additive genetic variance not mediated by ecological predictors. The last term corresponds to the non-inherited variance. We have

$$\begin{aligned} -2V_S \log \omega(z|\mathbf{x}^\star) &= (z - \bar{z}^\star)^2 \\ &= (\bar{z} - \bar{z}^\star + a\epsilon + e)^2 \\ &= (\bar{z} - \bar{z}^\star)^2 + 2(\bar{z} - \bar{z}^\star) \times (a\epsilon + e) + (a\epsilon + e)^2 \,. \end{aligned}$$

Thus, the expected value of the logarithm of $\omega(\mathbf{x}, \mathbf{x}^\star)$ (times $-2V_S$) is equal to

$$-2V_S \mathrm{E}[\log \omega(z|\mathbf{x}^\star)] = a^2 \mathcal{G}(\mathbf{x}, \mathbf{x}^\star) + a^2\sigma_\epsilon^2 + \sigma_e^2 \,.$$

### 3 Linear combination of predictors

In this section we assume that $\mathbf{x}$ represents a linear combination, $\mathbf{x} = \mathbf{x}'\mathbf{A}^T$, of causal variables $\mathbf{x}'$ for which traits are optimized. The coefficients of $\mathbf{A}$ are unknown. In this case, the geometric GO, which is computed from the non-causal predictors, is equivalent to a geometric GO computed from the causal variables

$$G^2(\mathbf{x}, \mathbf{x}^\star) = (\mathbf{x}' - \mathbf{x}'^\star)\mathbb{E}[\mathbf{c}^T\mathbf{c}](\mathbf{x}' - \mathbf{x}'^\star)^T \,,$$

where $\mathbf{c} = \mathbf{bA}$ are the effect sizes for the unobserved causal variables, $\mathbf{x}'$. This formula means that the definition of the geometric GO is robust to any correlation in the causal effects and works with unknown combinations of those effects. Because local stabilizing selection acts on traits controlled by causal variables, the geometric GO is proportional to the squared distance between causal environments in the ecological niche

$$G^2(\mathbf{x}, \mathbf{x}^\star) \propto (\mathbf{x}' - \mathbf{x}'^\star)\mathbf{C}^{-1}(\mathbf{x}' - \mathbf{x}'^\star)^T \,,$$

where $\mathbf{C}$ is the unknown covariance matrix of causal effects on fitness traits.

### 4 Relationships between GO statistics

This section provides details on the results presented in the main text paragraph "Unifying genetic offset statistics".

#### 4.1 Connection to environmental Euclidean distance

The geometric GO is determined by the eigenvalues of the covariance matrix of environmental effect sizes, $\mathbf{C_b} = \mathbb{E}[\mathbf{b}^T\mathbf{b}]$. If (and only if) the eigenvalues of the covariance matrix are equal, the geometric GO is proportional to the squared Euclidean distance between environmental predictors. According to the variational definition of eigenvalues and the spectral theorem, the quantity

$$G^2(\mathbf{x}, \mathbf{x}^\star) = (\mathbf{x} - \mathbf{x}^\star)\mathbf{C_b}(\mathbf{x} - \mathbf{x}^\star)^T \,,$$

is related to the Euclidean distance by the following inequalities

$$\lambda_d \|\mathbf{x} - \mathbf{x}^\star\|^2 \leq G^2(\mathbf{x}, \mathbf{x}^\star) \leq \lambda_1 \|\mathbf{x} - \mathbf{x}^\star\|^2 \,,$$

where $\lambda_d \leq \cdots \leq \lambda_1$ are the ranked eigenvalues of the covariance matrix, and $\|\mathbf{x}\|$ denotes the Euclidean norm of the environmental vector $\mathbf{x}$. If $\lambda_d = \lambda_1$, then the geometric GO is proportional to the squared Euclidean distance. If the geometric GO is proportional to the squared Euclidean distance (converse statement), the definition of eigenvalues as Raleigh quotients, which are all equal to the same value, implies that all eigenvalues are equal.

As a consequence of this result, the geometric GO is maximal in the direction of the first eigenvector of the covariance matrix, and it is minimal in the direction of the last eigenvector of the covariance matrix. For scaled environmental variables, the eigenvalues of the covariance matrix provide insights on the amount of information on local adaptation carried out by combinations of environmental predictors described by corresponding eigenvectors.

#### 4.2 Connection to the risk of nonadaptedness

The definition of the geometric GO shares similarities with the risk of non-adaptedness (Rona) proposed in (Rellstab *et al.*, 2016). The original definition of Rona is based on predictions of simple linear models adjusted for each environmental predictor considered separately. To remove bias due to population structure and to adopt a multivariate approach, Rona can be computed from predictions of a latent factor mixed model instead of a simple linear model (de Aquino *et al.*, 2022). With this modification, Rona writes as the average value of the difference between predicted allelic frequencies

$$\mathrm{Rona}(\mathbf{x}, \mathbf{x}^\star) = \mathbb{E}[|f(\mathbf{x}) - f(\mathbf{x}^\star)|] \,.$$

In the above equation, the latent factor terms cancel out, and we have

$$\mathrm{Rona}(\mathbf{x}, \mathbf{x}^\star) = \mathbb{E}[|(\mathbf{x} - \mathbf{x}^\star)\mathbf{b}^T|] \,,$$

where $\mathbf{b}$ represents the estimates of effect sizes obtained from the LFMM .

In comparison with Rona, the geometric GO considers squared differences instead of absolute differences. According to the Cauchy-Schwarz inequality, Rona is always lower than the squared root of the geometric GO,

$$\mathbb{E}[|f(\mathbf{x}) - f(\mathbf{x}^\star)|] \leq G(\mathbf{x}, \mathbf{x}^\star) = \sqrt{\mathbb{E}[(f(\mathbf{x}) - f(\mathbf{x}^\star))^2]} \,.$$

To go further in the comparison of Rona and the geometric GO, let us assume that the locus-specific environmental effect sizes are statistically independent and drawn from a $d$-dimensional Gaussian distribution at each locus

$$\mathbf{b}_\ell \sim N(0, \mathbf{C}_\mathbf{b}) \,, \quad \ell = 1, \ldots, L \,,$$

where $\mathbf{C_b}$ is identified to the theoretical covariance matrix. In empirical analysis, those assumptions may be satisfied when an LD pruning algorithm is applied to the genotype matrix prior to the GEA analysis, and when all loci are included in the computation of genomic offsets. According to the law of large numbers, we have

$$\frac{1}{L}\sum_{\ell=1}^{L} |(\mathbf{x}-\mathbf{x}^\star)\mathbf{b}_\ell^T| = \mathbb{E}[|f(\mathbf{x}) - f(\mathbf{x}^\star)|] \approx G(\mathbf{x}, \mathbf{x}^\star) \times \mathrm{E}[|Z|]\,,$$

where $\mathrm{E}[|Z|]$ is the expected value of a half-normal random variable. Under the Gaussian hypothesis, Rona is therefore expected to be proportional to the square root of the geometric GO. Note that the expected value, $\mathrm{E}[|Z|] = \sqrt{2/\pi}$, is less than one, so that the ranking of the two statistics shown in the inequality is preserved.

Next, to provide an explanation of Rona in a model of local stabilizing selection, we suppose that the local fitness gradient is described by an exponential (Laplace) curve. In this model, the fitness of the trait in environment $\mathbf{x}$ is defined as $\omega_{\mathrm{Laplace}}(z|\mathbf{x}) = \exp(-|z-\bar{z}|/2D_S)$, where $D_S$ is the width of the selection gradient ($D_S$ is used instead of $V_S$ to emphasize that the constant has the same dimension as a distance). Given $\mathbf{x}$ and $\mathbf{x}^\star$ and defining $\omega(\mathbf{x}, \mathbf{x}^\star)$ as the fitness value of an equilibrium trait in environment $\mathbf{x}$ being placed in the altered environment $\mathbf{x}^\star$, we have

$$-\log\omega(\mathbf{x}, \mathbf{x}^\star) = |\bar{z} - \bar{z}^\star|/2D_S = |a(\mathbf{x}^\star - \mathbf{x})\mathbf{b}^T|/2D_S\,.$$

Assuming $\mathbf{b} \sim N(0, \mathbf{C_b})$, the distribution of the absolute difference describes as

$$|\bar{z}^\star - \bar{z}| \sim |a|\,|Z|\,\mathcal{G}(\mathbf{x}, \mathbf{x}^\star).$$

where $|Z|$ has half-normal distribution. According to the above equation, the expected value of the logarithm of altered fitness varies in proportion with Rona

$$-\mathrm{E}[\log \omega(\mathbf{x}, \mathbf{x}^\star)] \approx \mathrm{Rona}(\mathbf{x}, \mathbf{x}^\star)/2D_s\,,$$

where $1/D_s = (|a|\sqrt{2/\pi})/D_S$. Like the geometric GO, Rona estimates the logarithm of fitness in the altered environment, but does it for an exponential selection gradient instead of a Gaussian gradient. The ability of each statistic to predict fitness in the altered environment will thus be linked to the shape of the selection gradient.

#### 4.3 Connection to redundancy analysis (RDA)

The definition of the geometric GO (genetic gap) as a distance in environmental space is closely related to RDA GO introduced in (Capblancq *et al.*, 2020; Capblancq and Forester, 2021). RDA first adjusts $d$ linear predictors to the genotype matrix by using linear regression, and then performs a principal component analysis (PCA) of the fitted values. With a modification ignored below, Capblancq *et al.*'s genetic offset was defined as the Euclidean distance between projections of current and modified environmental predictors on the first principal components of the fitted values. Here we assume that 1) RDA is performed by using $d$ environmental predictors and the $K$ latent factors of the GEA study, 2) Projections in RDA space include all $(d + K)$ RDA components, i.e., as many components as the dimension of environmental and latent space. Under these conditions, this section provides a mathematical proof that the squared values of the RDA GO is equivalent to the geometric GO. This property results from a general property of PCA projections which preserve Euclidean distance.

Under the first hypothesis, the linear predictors are defined in the matrix $\mathbf{X}\mathbf{B}^T + \mathbf{U}\mathbf{V}^T$ where $\mathbf{X}$ is the $n \times d$ design matrix, representing $n$ samples of $d$-dimensional environmental predictors, $\mathbf{B}$ is the $L \times d$ matrix of effect sizes, $\mathbf{U}$ is the $n \times K$ factor matrix, $\mathbf{V}$ is the $L \times K$ matrix of factor loadings, and $L$ is the number of genomic loci. PCA decomposes the predicted values as

$$(\mathbf{X}\mathbf{B}^T + \mathbf{U}\mathbf{V}^T)/\sqrt{n-1} = \mathbf{Q}\mathbf{P}^T \,,$$

where $\mathbf{Q}$ are the PC scores (or sample projections), and $\mathbf{P}$ are the corresponding loadings (of dimension $L \times n$). For any $d$-dimensional vector of environmental predictors, $\mathbf{x}$, and $K$-dimensional factor, $\mathbf{u}$, the projection in the RDA space is given by

$$\mathbf{proj}(\mathbf{x}) = (\mathbf{x}\mathbf{B}^T + \mathbf{u}\mathbf{V}^T)\mathbf{P} \,.$$

Often, $\mathbf{x}$ and $\mathbf{u}$ correspond to values measured at some sample location, but the projection above could be computed at any arbitrary location for which $\mathbf{x}$ (and $\mathbf{u}$) are given. For a modified environment, $\mathbf{x}^\star$, at the same location as $\mathbf{x}$, we have

$$\mathbf{proj}(\mathbf{x}^\star) = (\mathbf{x}^\star\mathbf{B}^T + \mathbf{u}\mathbf{V}^T)\mathbf{P} \,.$$

Since $\mathbf{P}$ is a linear transformation, the $\mathbf{u}\mathbf{V}^T$ term disappears when differentiating, leading to

$$\mathbf{proj}(\mathbf{x}^\star) - \mathbf{proj}(\mathbf{x}) = (\mathbf{x}^\star - \mathbf{x})\mathbf{B}^T\mathbf{P}\,.$$

The squared Euclidean distance in the RDA space can be computed as follows

$$\|\mathbf{proj}(\mathbf{x}^\star) - \mathbf{proj}(\mathbf{x})\|^2 = (\mathbf{x}^\star - \mathbf{x})\mathbf{B}^T\mathbf{P}\mathbf{P}^T\mathbf{B}(\mathbf{x}^\star - \mathbf{x})^T\,.$$

A property of PCA is that $\mathbf{P}$ is a semi-orthogonal matrix, such that

$$\mathbf{P}\mathbf{P}^T = \mathbf{I}\,,$$

where $\mathbf{I}$ is the $L \times L$ identity matrix. In addition, we have

$$\mathbf{C_b} = \mathbb{E}[\mathbf{b}^T\mathbf{b}] = \mathbf{B}^T\mathbf{B}/L\,.$$

Thus, we have

$$\frac{1}{L} \times \|\mathbf{proj}(\mathbf{x}^\star) - \mathbf{proj}(\mathbf{x})\|^2 = (\mathbf{x}^\star - \mathbf{x})\mathbf{C_b}(\mathbf{x}^\star - \mathbf{x})^T = G^2(\mathbf{x}, \mathbf{x}^\star)\,.$$

In other words, the geometric GO is equal to the squared Euclidean distance in RDA space times the inverse of the number of loci considered in the analysis.

#### 4.4 Connection to gradient forest genetic offset

Gradient forest (GF) is a nonlinear and nonparametric approach based on random forests that does not assume any a priori model of selection, but infers it from the observed data (Ellis *et al.*, 2012; Fitzpatrick and Keller, 2015). The GF genetic offset is built from importance curves ($\mathrm{IC}_j$), estimated for each of the $d$ environmental predictors as follows

$$\mathrm{GF}^2(\mathbf{x}, \mathbf{x}^\star) = \sum_{j=1}^{d} .(\mathrm{IC}_j(x_j) - \mathrm{IC}_j(x_j^\star))^2 \,.$$

The construction of the GF GO statistic shares similarities with the geometric GO and with the RDA GO, in which projections are weighted by the eigenvalues of the covariance matrix, representing the importance of linear combinations of environmental predictors inferred from the data

$$G^2(\mathbf{x}, \mathbf{x}^\star) = \frac{1}{L} \sum_{j=1}^{d} (\mathrm{proj}_j(\mathbf{x}) - \mathrm{proj}_j(\mathbf{x}^\star))^2 \,.$$

An important difference between the geometric GO and the GF GO is linearity versus nonlinearity. Nonlinearity makes the GF model more flexible than linear models, but the additional degrees of freedom may not always be desirable when resolving bias-variance trade-offs.

### 5 Genetic offset computations

#### 5.1 geometric GO

Computations of the geometric GO (genetic gap) were performed with the function `genetic.gap` in the `R` package LEA version 3.9.5 (Gain and François, 2021).

#### 5.2 RDA genetic offset

We used redundancy analysis (RDA) to fit a simple linear model to the genotype matrix and computed the genetic offset as the squared distance between the projections of environmental predictors on the principal components of the fitted values. To correct for the confounding effect of demography, we included the $K$ factors estimated in the GEA analysis as predictors in the RDA linear model. Our implementation differs from the implementation of Capblancq *et al.* (2020), that used projections along two axes (RDA subspace) and incorporated weightings for each dimension. For consistency with the geometric approach, we reported the squared values of Euclidean distance in the full RDA space. Our implementation made the RDA GO theoretically equivalent to the geometric GO.

#### 5.3 Gradient forest genetic offset

We used gradient forests (GF) to compute genetic offsets that model the allelic composition of populations by using nonlinear functions of environmental gradients, called cumulative importance curves (Ellis *et al.*, 2012; Fitzpatrick and Keller, 2015). In GF, genetic offset statistics were computed as squared differences of the cumulative importance values for two sets of environmental predictors. Computations were performed using the `R` package `gradientForest` version 0.1. To correct for population structure, we first adjusted the genotype matrix on the $K$ factors estimated in the GEA analysis, and performed GF analyzes on the residuals of the linear model. For consistency with the geometric approach, we reported squared values for the genetic offset.

#### 5.4 Risk of nonadaptedness

Following (Rellstab *et al.*, 2016), the risk of nonadaptedness (Rona) was computed as the average value of the absolute distance between fitted and predicted allelic frequencies across genomic loci. In Rona, allelic frequencies were adjusted on environmental predictors using simple linear models. To remove bias due to population structure, Rona was computed from predictions of a latent factor mixed model instead of a simple linear model (de Aquino *et al.*, 2022). Allelic frequencies predicted by observed environnement were used instead of observed allelic frequencies.

#### 5.5 Variational autoencoder genetic offset

We used genomic and environmental data in a deep learning generative model to establish a statistical relationship between environmental factors and allelic frequencies. We used this relationship to simulate the required genomic composition to be adapted to altered environmental conditions. This approach provides constrained predictions of allelic frequencies, $f_c(\mathbf{x})$, and avoids their linear approximations. Following (Kingma and Welling, 2013; Liao and Lin, 2021), we implemented a conditional variational auto-encoder that consisted of two probabilistic encoders with a teacher-student relationship. We estimated a genetic offset from the generative model by using the formula $\mathbb{E}[(f_c(\mathbf{x}) - f_c(\mathbf{x}^\star))^2]$, where $f_c(\mathbf{x})$ and $f_c(\mathbf{x}^\star)$ were computed from the deep learning generative model. The objective of this implementation was to check whether the results were consistent with the geometric GO or not. To do this, we plotted the constrainted genetic offset values against the geometric GO values in four case studies (see extended simulation study).

### 6 Evaluating the bias of linear predictors

To evaluate the impact of the approximation of allelic frequencies by linear functions, we introduced the logit link, $\sigma(x) = 1/(1+e^{-x})$, in a logistic regression model of allelic frequencies on environmental predictors, $\mathbf{x}$, and latent factors, $\mathbf{u}$. After adjusting the model to the data, predicted allelic frequencies took the following form

$$f_c(\mathbf{x}) = \sigma(\mathbf{x}\mathbf{a}^T + \mathbf{u}\mathbf{w}^T)\,,$$

where $\mathbf{a}$ and $\mathbf{w}$ are the effect sizes and loadings estimated in the logistic regression model. Those constrained predictors take their value between 0 and 1 and have direct interpretation as probabilistic quantities.

For small local environmental change, a Taylor expansion of the difference between predicted allelic frequency led to

$$f_c(\mathbf{x}^\star) - f_c(\mathbf{x}) \approx f_c(\mathbf{x})(1 - f_c(\mathbf{x}))(\mathbf{x}^\star - \mathbf{x})\mathbf{a}^T\,.$$

This can be identified with the equation for the linear predictor,

$$f(\mathbf{x}^\star) - f(\mathbf{x}) = (\mathbf{x}^\star - \mathbf{x})\mathbf{b}^T\,,$$

by setting

$$\mathbf{b} = f_c(\mathbf{x})(1 - f_c(\mathbf{x})) \times \mathbf{a} = H_0 \times \mathbf{a}/2\,,$$

where $H_0$ is the genetic heterozygosity at the locus considered. For small environmental changes, the genetic distance computed from the constrained predictors $\mathbb{E}[(f_c(\mathbf{x}^\star) - f_c(\mathbf{x}))^2]$ is then expected to approximate the geometric GO accurately.

### 7 Extended simulation study

This section details the settings for the simulations described in the main text paragraph “Extended simulation study”. Spatially-explicit individual-based simulations were performed using SLiM 3.7 (Haller and Messer, 2019). Each individual genome contained neutral mutations and quantitative trait loci (QTLs) under stabilizing selection controlled by two causal environmental variables. The viability of an individual genome, defined as the probability of survival in the next generation, was computed as the product of density regulation and fitness. We designed four classes of scenarios, including weakly or highly polygenic QTLs, and weak or high correlation of causal environmental variables with the first principal components of the genotype matrix (population structure). In scenarios with high polygenicity, traits controlled by 120 mutations with additive effects were matched to each environmental variable by stabilizing selection. In weakly polygenic scenarios, the traits were controlled by 10 mutations. Scenarios with high confounding effects were initiated in a demographic range expansion process, creating correlation between environmental gradients and allelic frequencies at the genome level. For each scenario, thirty replicates were run with distinct seed values of the random generator. At the end of a simulation, individual geographic coordinates, environmental variables before and after environmental change, and individual fitness values before and after abrupt environmental change were recorded.

#### 7.1 Simulation details

**Geography and environmental gradients.** All simulations took place in a two-dimensional square area of size 10 units. A first environmental gradient, $x_1$, varied from west to east, and a second environmental gradient, $x_2$, varied from south to north. Those environmental gradients influenced the viability of individual genomes, and no other variable was causal for selection in the simulation.

**Life cycle and evolutionary rates.** The timescale of simulated evolution was arbitrarily chosen so that local adaptation could take place in reasonable time. A generation was considered as a succession of lifetime periods with reproduction, migration, and selection. The number of periods in an individual lifetime did not result in any change in the genotype matrix at the end of the simulation (only timescale was modified). The scales of mutation, recombination, migration, reproduction rates and the ratio of autogamy to allogamy were thus expressed in relative units. The numbers below were given for a generation time corresponding to a single lifetime period. Neutral mutations appeared at a rate of $3.0 \times 10^{-6}$ per base pair per generation. The recombination rate was equal to $1.0 \times 10^{-2}$ per base pair per generation. The genome size was around 110 kb. Individuals migrated according to a random walk of step taken in (-0.1, 0.1) on each geographic axis. At the end of a generation, an individual genome was transmitted to the next generation with probability equal to the product of density regulation and local adaptation (see below). An additional number of offspring was drawn from a Poisson distribution of parameter 10%. A random mate was chosen in a radius of size 0.5 units. The birth location corresponded to the barycenter of the parent locations plus a Gaussian noise of standard deviation $\sigma = 0.05$.

**Phenotypes.** An individual genome contained selectively neutral mutations as well as QTLs controlled by non-neutral mutations. Two QTLs were simulated, each underlying a specific phenotypic trait under selection controlled by one environmental variable. In each QTL, the value of the trait was determined by additive effects. A derived allele in a QTL increased the value of the trait by a fixed amount which varied according to the level of polygenicity in each scenario.

**Density regulation.** The density of individuals was regulated by spatial competition according to the following formula for the survival of an individual in the next generation (Haller and Messer, 2019)

$$\text{Survival} = \frac{11\pi S^2}{N_{\text{ind}} + 1} \tag{1}$$

where $N_{\text{ind}}$ was the number of individuals within a circle of radius $S = 0.8$ around each individual.

**Fitness.** Fitness was computed according to the formula for Gaussian stabilizing selection (Burger, 2000)

$$\text{Fitness} = \exp\left(-\frac{1}{2}(\mathbf{z} - \bar{\mathbf{z}})C^{-1}(\mathbf{z} - \bar{\mathbf{z}})^T\right) \tag{2}$$

where $\bar{\mathbf{z}} \approx \mathbf{x}$ is the vector of environmental variables at the geographic coordinates of the individual, $\mathbf{z}$ is the vector of phenotypic traits and $C$ is a covariance matrix for selection coefficients. We used

$$C = \begin{pmatrix} 0.04 & 0 \\ 0 & 0.04 \end{pmatrix} .$$

The value of 0.04 was chosen in order to ensure that a local adaptation equilibrium was reached after a reasonable simulation time, before environmental change occurred.

**Evolutionary phases.** Each simulation consisted of two distinct phases. The first phase was a purely demographic phase shaping population structure, and the second phase allowed populations to evolve traits until they reached an equilibrium for local adaptation. During the demographic phase, mutations accumulated to create genetic diversity in the population. At the beginning of the adaptive phase, selection on QTLs started from standing variation in a soft sweep, with an initial frequency of around 300 for each derived allele. The initial frequency of 300 was chosen so that derived alleles were not removed by genetic drift in a fluctuating population size of around 2,000 individuals. At the end of the adaptive phase, the adaptive traits matched the values of the causal environmental variables locally. Environmental changes consisted of instantaneous and heterogeneous alterations of the causal environmental variables. This was obtained by adding local values randomly drawn in the range (-.3, +.3) to each causal variable. In altered local environments, the values of fitness were computed at the time the change occurred.

#### 7.2 Types of simulation scenarios

We designed four classes of scenarios, including weakly or highly polygenic QTLs, and weak or high correlation of environmental gradients with population structure.

**Weakly and highly polygenic traits.** We varied the number of mutations that controlled the two QTLs underlying the traits involved in the process of local adaptation. In scenarios with high polygenicity, the traits were controlled by 120 mutations with additive effects. The derived allele of a mutation increased the value of a trait by 0.005, allowing each trait to vary between 0 and 1 in order to match their environment ranging in the same interval. In weakly polygenic scenarios, the traits were controlled by 10 mutations. A derived allele of a mutation increased the value of the trait by 0.05, allowing each trait to vary between 0 and 1.

**Weak and high correlation with population structure.** We designed scenarios displaying weak or high correlation of environmental gradients with population structure. The difference between scenarios was in the demographic phase. In low correlation scenarios, individual migrated spatially according to a random walk with steps in the interval (-0.3, 0.3), allowing high levels of gene flow between geographic compartments. In high correlation scenarios, the steps were taken in (-0.1, 0.1) and levels of gene flow were much lower. At the start of the simulation, the northern part of the map was hostile to individuals, and this area was made gradually suitable for individuals along generations. By simulating range expansion from the south, population structure was oriented along the second environmental variable.

##### 7.3 Data used in "Validation of the theory".

To illustrate the theory, we considered an example in which $n = 380$ individuals were sampled from the simulated range. In this example, the geometric GO was computed to check whether it correlates linearly with the logarithm of fitness after environmental change, as predicted by the quantitative theory. Four predictors were measured for each individual. The first two predictors, $x_1$ and $x_2$, was causally involved in the evolutionary process of local adaptation by influencing allelic frequencies at a subset of adaptive loci. Two additional predictors, $x_3$ and $x_4$, were included, defined as linear combinations of the first two predictors. The third variable, $x_3$, exhibited high correlation with $x_1$ ($r = 0.71$) and with $x_2$ ($r = 0.71$). The fourth variable, $x_4$, was also highly correlated with $x_1$ ($r = 0.93$) and $x_2$ ($r = -0.35$). The median values of $x_1$ and $x_2$ determined four broad types of environments, termed *dry/warm* to *wet/cold* conditions, by analogy with bioclimatic conditions. In this simulation, the correlation between the first principal axis of the genotype matrix and $x_2$ was equal to $-0.93$, and the correlation between the second principal axis and $x_1$ was equal to 0.88.

#### 8 Code and data availability

The codes necessary to reproduce the simulations and data analyses of this study are available at `https://github.com/bcm-uga/geneticgap` under GNU

General Public License v3.0.
