## Supplementary Figures and Tables for "A quantitative theory for genomic offset statistics"

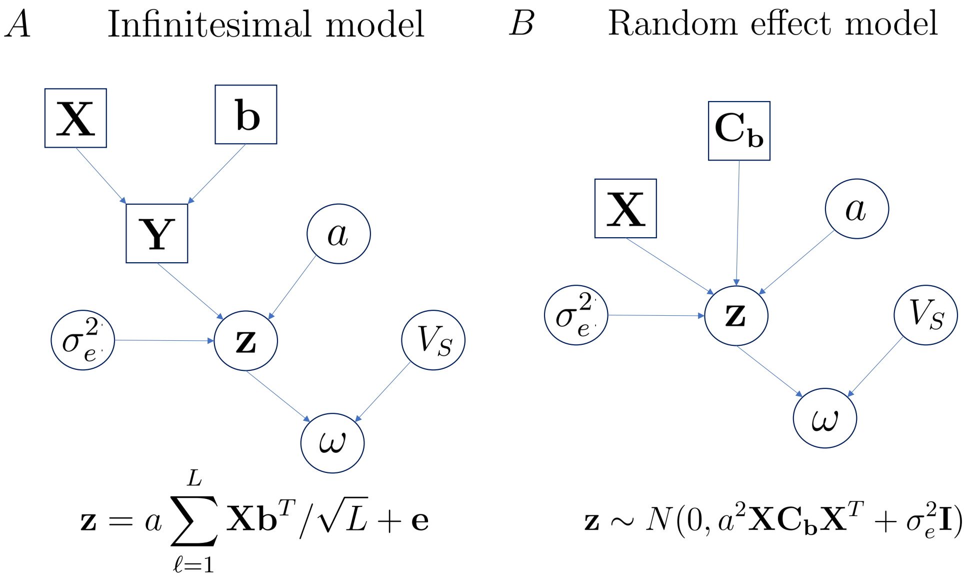


**Figure S1. Directed acyclic graph for fitness traits.** A) Infinitesimal model in which allelic effects are predicted from a GEA model. B) Equivalent random effect model in which the covariance matrix of random effects is inferred from the GEA model. Confounding factors and GEA residuals were omitted for simplification. Data and parameters: matrix of environmental predictors, $\mathbf{X}$, matrix of allelic frequencies, $\mathbf{Y}$, matrix of environmental effect sizes, $\mathbf{b}$, covariance of effect sizes, $\mathbf{C_b}$, vector of unobserved traits, $\mathbf{z}$, fitness function, $\omega$, genetic variance, $a^2$, width (variance) of the selection gradient, $V_S$, non-inherited variance, $\sigma_e^2$, number of genetic loci, $L$. Squares represent observed variables, circles represent hidden variables.

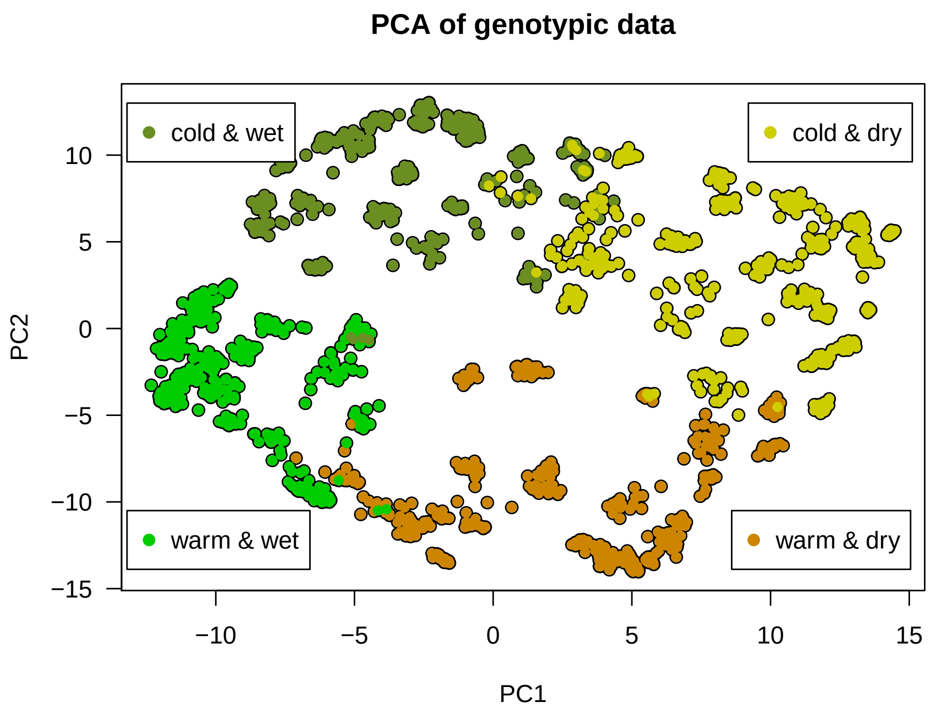


**Figure S2. Simulated example data set: PCA of the genotype matrix.** The projections of samples in the principal component space are colored according to their membership in one of the four quadrants of the two-dimensional environmental space (temperature and precipitation). For example, 'warm and wet' corresponds to nonnegative temperatures and precipitations. All variables are expressed as deviation from the population mean.

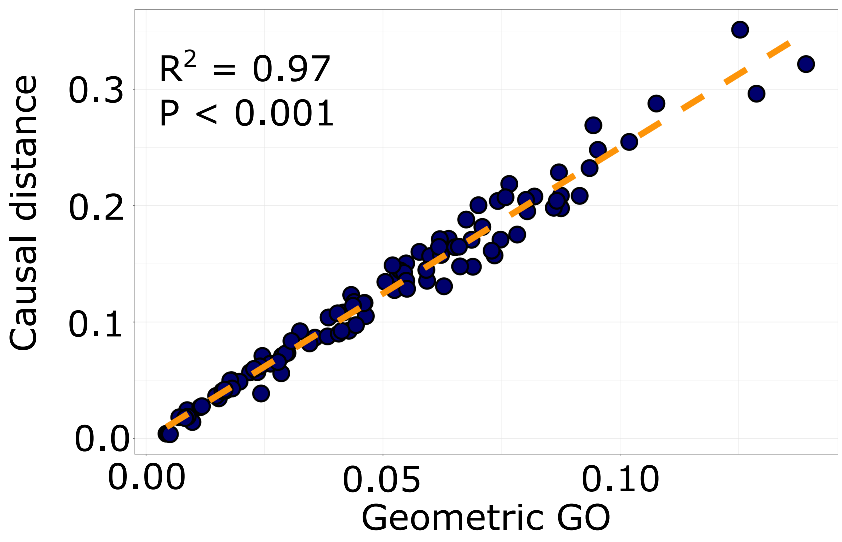


**Figure S3. Simulated example data set: Geometric GO and causal quadratic distance.** Geometric offset as a function of the squared distance between environment that determined the intensity of the Gaussian stabilizing selection in the simulation.

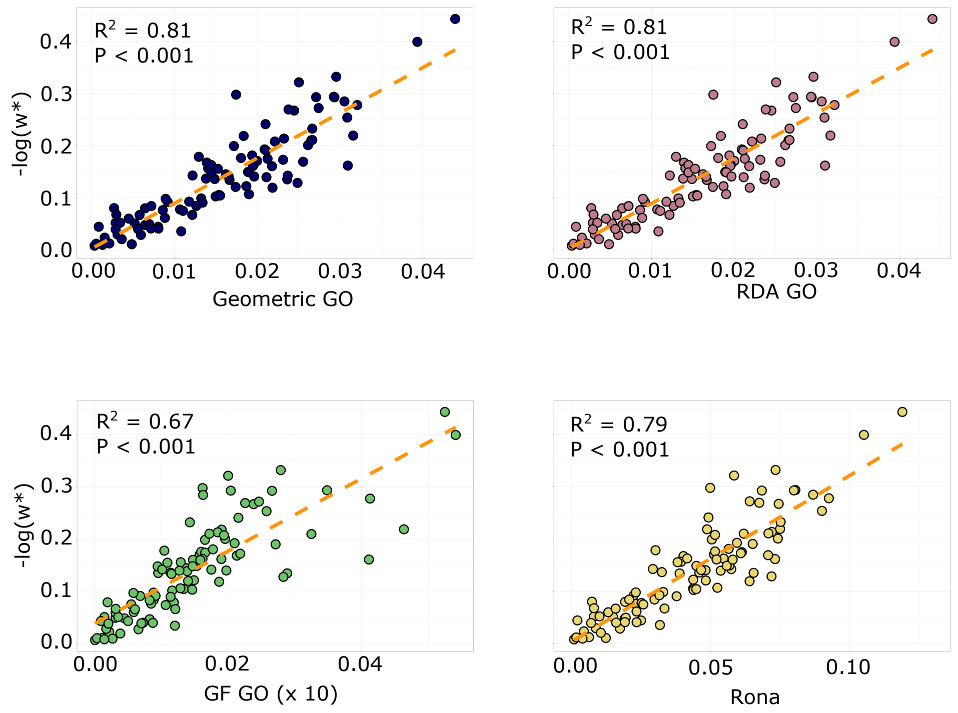


**Figure S4. Extended simulation study. Regression plots.** Typical regression plots for simulated data with weakly polygenic fitness traits and high levels of correlation between population structure and environment. The geometric GO, RDA GO, Rona and GF GO included corrections based on latent factors estimated in the GEA. GF GO values were multiplied by a factor of ten.

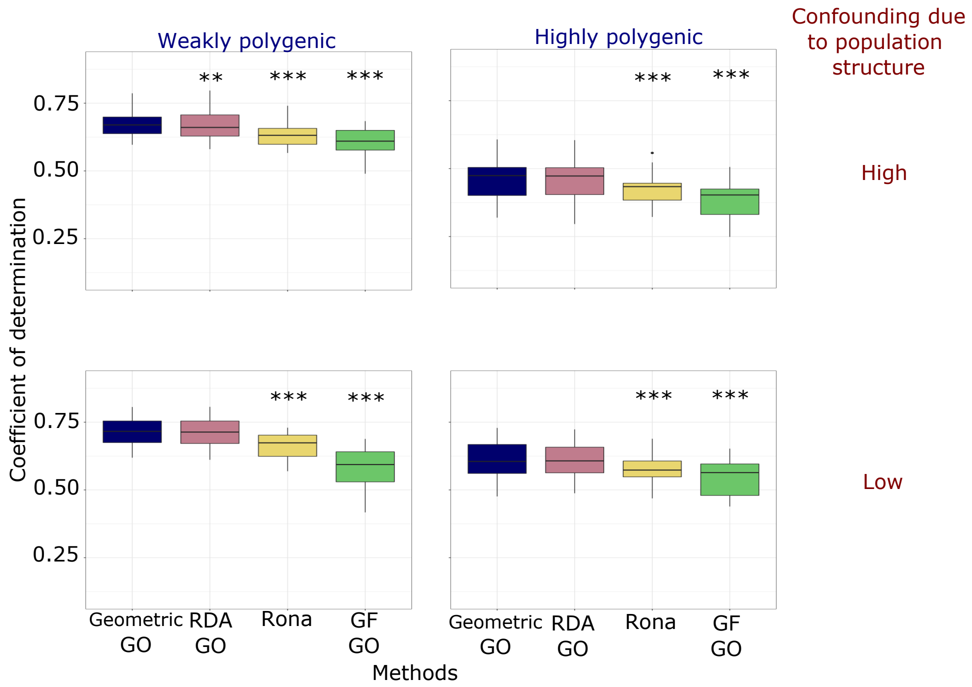


**Figure S5. Simulation study. Relative performances of uncorrected GO statistics.** Predictive ability of GO statistics without corrections using latent factors, following the original definitions for RDA GO, Rona and GF GO. The performances are lower than with corrections using latent factors (see Figure 3). Significance values were based on paired t-tests of the difference in mean performance for each GO statistic relative to the geometric GO. Significance code: *** $P < 0.001$, ** $< 0.01$.

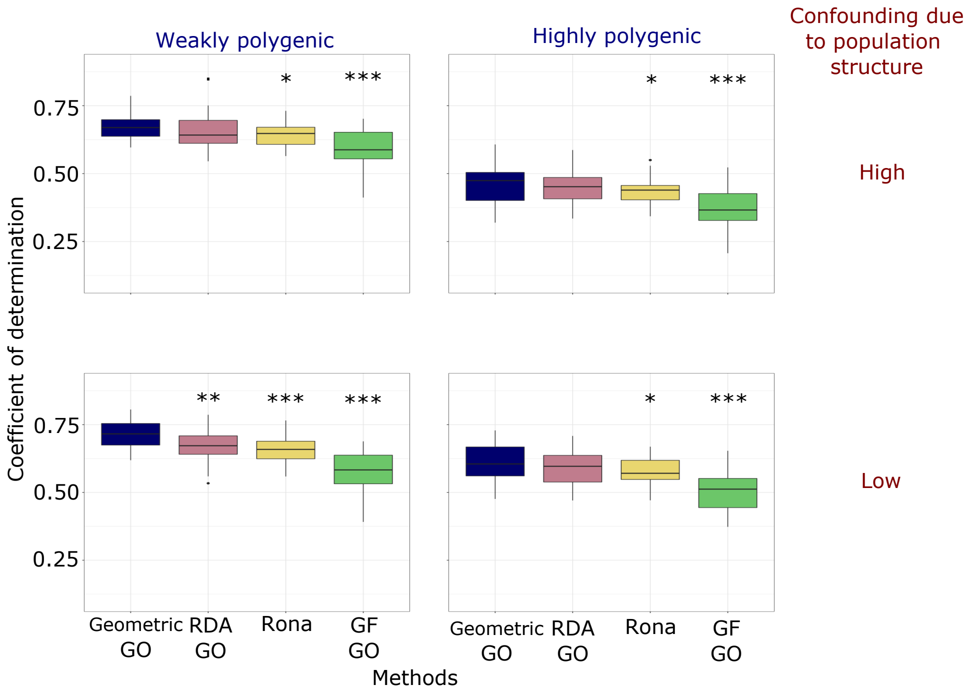


**Figure S6. Simulation study. Relative performances of GO statistics with corrections for population structure.** Predictive ability of GO statistics using correction based on principal components of the genotype matrix (10 PCs representing population structure). The performances are lower than with corrections using latent factors (see Figure 3). Significance values were based on paired $t$-tests of the difference in mean performance for each GO statistic relative to the geometric GO. Significance code: *** $P < 0.001$, ** $< 0.01$, * $< 0.05$, . $< 0.10$.

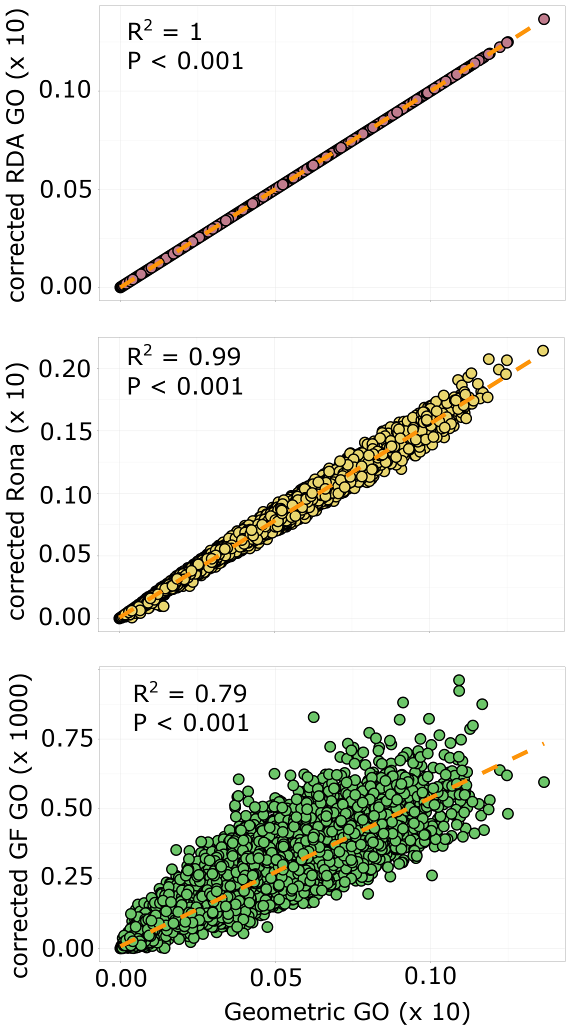


**Figure S7. Simulation study. GO statistic as function of the geometric GO.** Simulation values for the RDA GO, Rona, and the GF GO are plotted against the values of the geometric GO. GO statistics were computed on all loci included in the genotype matrix for 30 replicates of four scenarios (all results represented simultaneously).

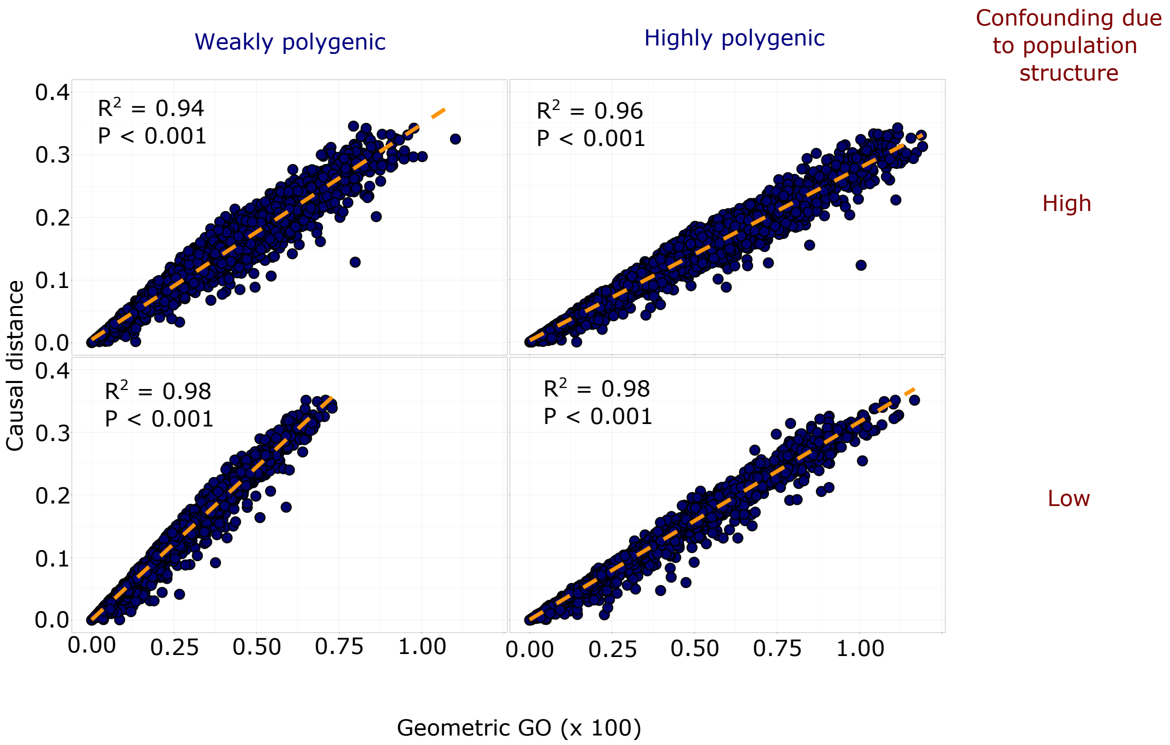


**Figure S8. Simulation study. Correlation with causal quadratic distance.** Squared distance between the variables that causally impact fitness in simulations and their altered values as a function of the geometric GO computed on all loci. High levels of correlation indicate that predictions made with the geometric GO are near-optimal (all results represented simultaneously, 30 replicates).

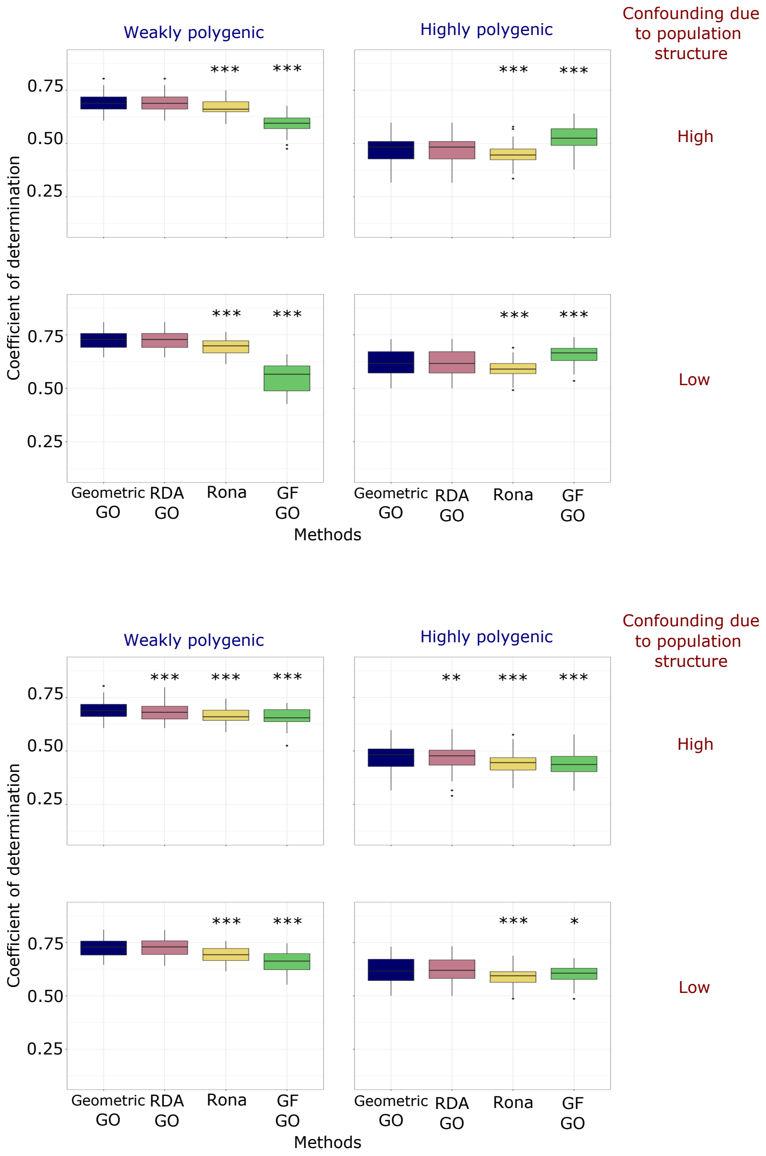


**Figure S9. Simulation study. Performances of GO based on all loci.** Ability of GO statistics to predict fitness calculated on the basis of all genomic data included in the genotype matrix. GO statistics were calculated with corrections using 10 latent factors (top panel), and without those corrections (bottom panel). Predictive performances were slightly lower than with subsets of genomic loci identified in GEA analyzes (Figure 3). Significance values were based on paired t-tests of the difference in mean performance for each GO statistic relative to the geometric GO. Significance code: *** $P < 0.001$, ** $< 0.01$, * $< 0.05$, . $< 0.10$.

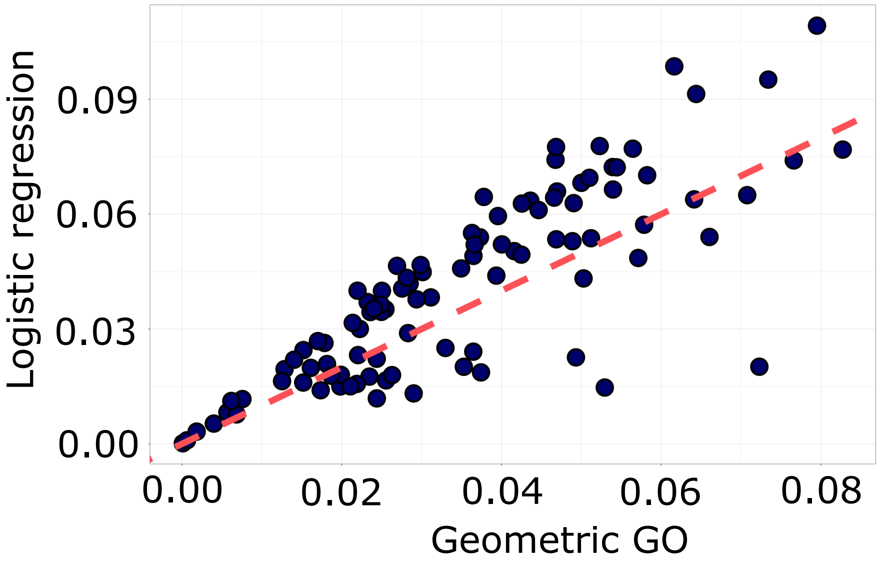


**Figure S10. Bias of linear models.** Accuracy of approximation of the quadratic distance between constrained predictors, $\mathbb{E}[(f_c(\mathbf{x}) - f_c(\mathbf{x}^\star))^2]$, based on logistic regression, as a function of the geometric GO, which is based on unconstrained linear predictors.

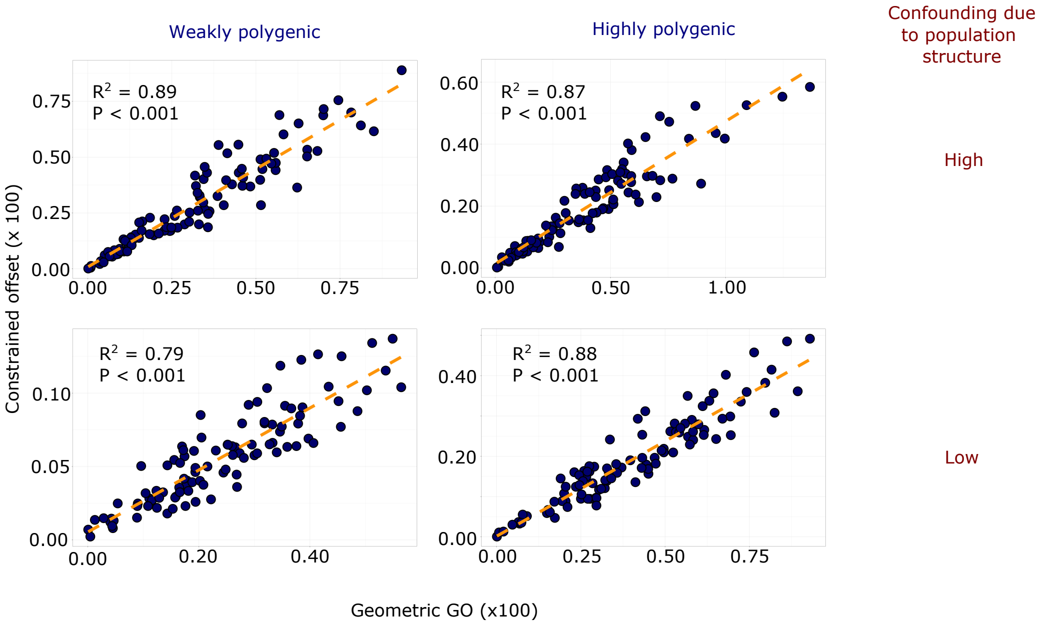


**Figure S11. Geometric GO computed from a machine learning model.** Accuracy of approximation of the quadratic distance between constrained allelic frequency predictors, $\mathbb{E}[(f_c(\mathbf{x}) - f_c(\mathbf{x}^\star))^2]$, based on conditional variational autoencoders, as a function of the geometric GO, which is based on linear regression models. GO computations were based on all loci included in the genotype matrix. The data analyzed correspond to the simulation scenarios considered in the extended simulation study.

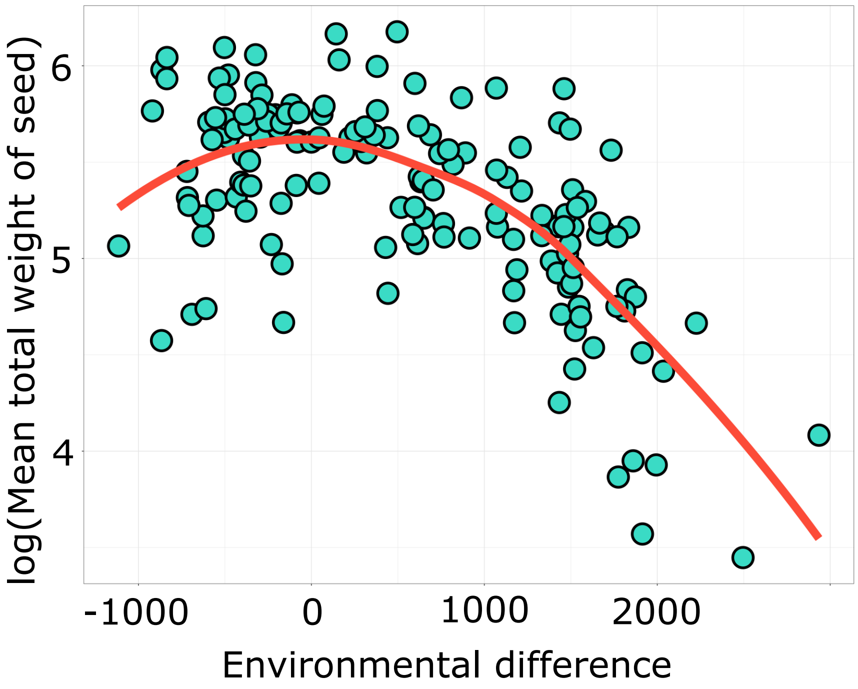


**Figure S12. Selection gradient in the pearl millet common garden experiment.** Logarithm of fitness values as a function of environmental predictors projected on their first PCA axis. The values were centered to represent the difference in environment between the landrace origin and the common garden location in Niger. Fitness was evaluated as the mean total seed weight for each pearl millet landrace. The fitted local regression curve shows a good agreement with a quadratic selection gradient.

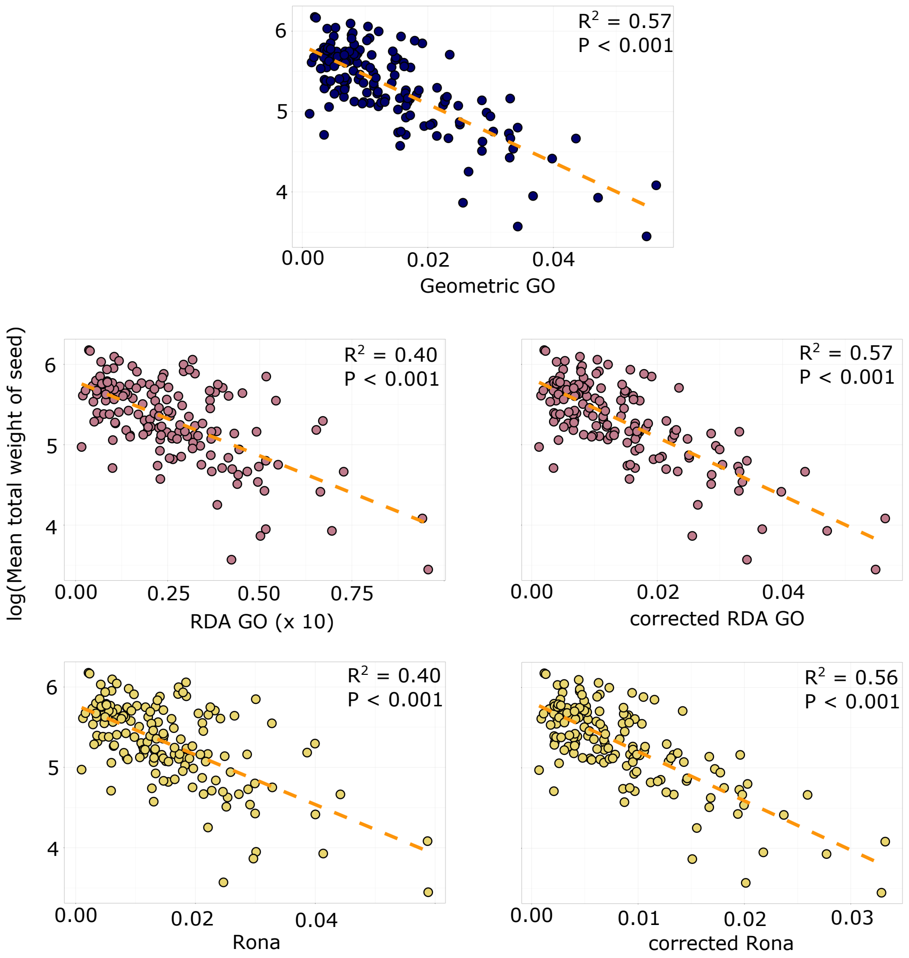


**Figure S13. Logarithm of fitness in the common garden as a function of GO statistics (whole genome).** Statistics were computed by using environmental effect sizes estimated from a GEA model at all genomic loci in the genotype matrix. For the RDA GO and for Rona, effect sizes were computed without correction for confounding factors (left column) or with correction for confounding factors using 10 LFMM factors (right column). Fitness was evaluated as the mean total weight of seeds for each of the 170 pearl millet landraces. Predictive performances were lower than with subsets of genomic loci identified in GEA analyzes and including latent factors improved the performances of methods (Figure 5).

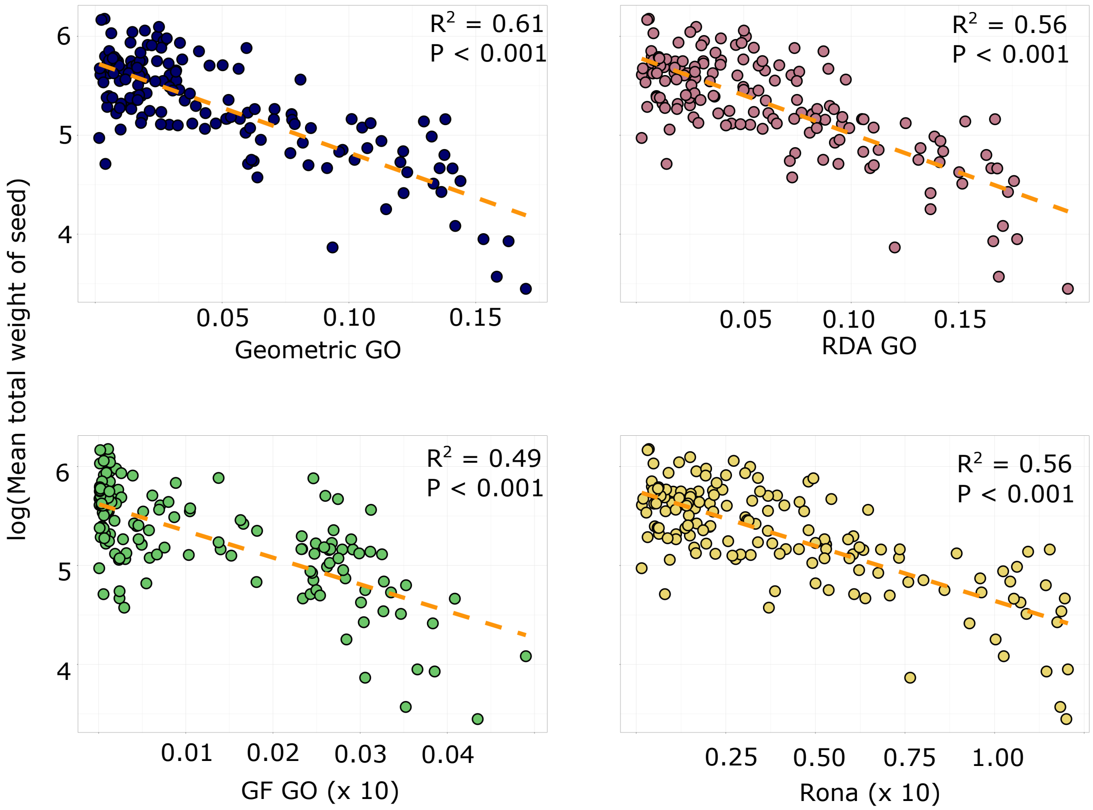


**Figure S14. Logarithm of fitness in the common garden as a function of GO statistics (uncorrected GO).** For RDA, Rona and GF, effect sizes were computed without correction for confounding factors. GO statistics were computed by using environmental effect sizes at loci detected from the GEA model. Fitness was evaluated as the mean total weight of seeds for each of the 170 pearl millet landraces. Latent factors improved the performances of methods (Figure 5).

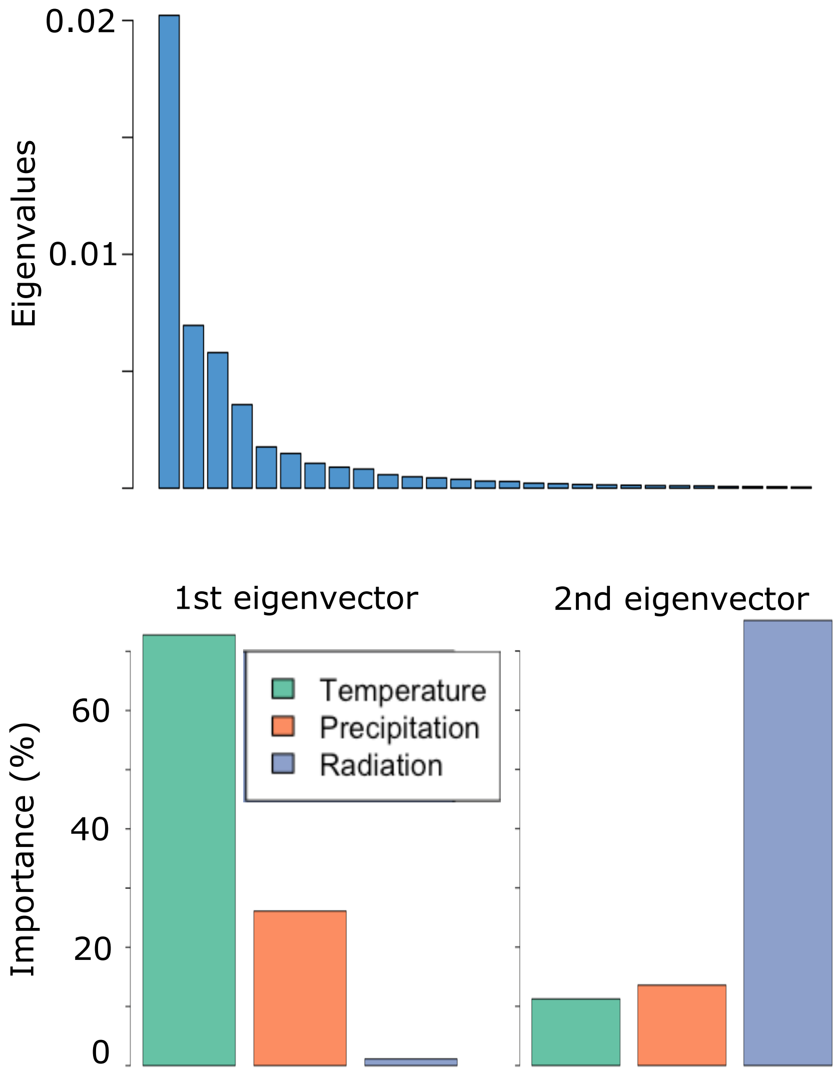


**Figure S15. Relative importance of bioclimatic categories.** Top: Eigenvalues of the covariance matrix of environmental effect sizes, showing that three axes explain most of the variation. Bottom: Relative importance of environmental predictors in driving the first axes. Relative importance was computed based on the squared loadings on the first and second axes.

**Table S1. Pearl millet analysis**. $J$ statistics and significance values for comparison of squared correlations between the geometric GO and the other GO statistics ($n = 170$).

| | **uncorrected Rona** | **corrected Rona** | **uncorrected RDA GO** | **uncorrected GF GO** | **corrected GF GO** |
|---|---|---|---|---|---|
| $J$ | 4.9 | 1.7 | 4.2 | 6.6 | 7.2 |
| $P$ | $2.5 \times 10^{-6}$ | 0.08 | $4.1 \times 10^{-5}$ | $6.5 \times 10^{-10}$ | $2.2 \times 10^{-11}$ |

GO statistics were computed using loci detected in the GEA study (FDR level = 10%).

**Note:** The $J$-test compares the fit of two regression models with distinct predictors. To compare model fit, the fitted values of model 1 are included into model 2 and vice versa. The $J$-test is the marginal test of the fitted values in the augmented model.
